# Stress-induced ribosome degradation in *Bacillus subtilis* is mediated by the RNase Y-specificity complex

**DOI:** 10.1101/2025.06.27.661978

**Authors:** Fabián A. Cornejo, Kristina Driller, Rina Ahmed-Begrich, Katja Schmidt, Michael Jahn, Vivekanandan Shanmuganathan, Karin Hahnke, Florian Kondrot, Thomas F. Wulff, Sebastian Rämisch, Kathirvel Alagesan, Emmanuelle Charpentier, Kürşad Turgay

## Abstract

Limiting the synthesis and activity of ribosomes is crucial for adaptation to stresses, such as heat or nutrient starvation. In *Bacillus subtilis,* this can be achieved through the coordinated action of the alarmones (p)ppGpp and the transcription factor Spx. Here, we performed a genetic screen to uncover novel factors contributing to the heat shock response of *B. subtilis*. We identified the Y-complex, which confers specificity to the endonuclease RNase Y, as a critical player during stress conditions, such as heat or transition into stationary phase. This protein complex is required for processing diverse RNAs, notably the maturation of mRNAs encoding proteins involved in translation and metabolism. We further demonstrate that the Y-complex and RNase Y initiate the degradation of rRNAs of mature ribosomes, lowering their abundance. We propose that the Y-complex is a regulatory hub that modulates gene expression, adjusts protein synthesis and resource allocation.

## Introduction

Bacteria can sense, adapt to, and survive stressful environmental changes facilitated by various general and specific stress response systems. Proteotoxic stresses, like high temperatures, significantly disrupt protein folding and can lead to the formation of toxic protein aggregates. Chaperones and proteases are expressed as a response due to the inactivation of the transcriptional repressors HrcA and CtsR in *Bacillus subtilis*^1–4^. The activation of the protein quality control (PQC) allows the refolding of un-or misfolded proteins by different chaperone systems or the degradation of irreparable proteins through AAA+ protease complexes like ClpCP^5,6^.

In addition to the repairing or removal of protein aggregates, inhibiting translation is crucial for survival during proteotoxic stresses, as it can alleviate the extra load on the PQC system^7–10^. In *Bacillus subtilis*, the nucleotide second messengers guanosine tetraphosphate and guanosine pentaphosphate, collectively referred to as (p)ppGpp, and the transcription factor Spx independently regulate synthesis and activity of ribosomes during stress^7,11^. Spx is a redox-sensitive heat shock transcription factor that controls the expression of redox chaperones and simultaneously represses transcription of translation-related genes during oxidative and heat shock response^11–14^.

The alarmones (p)ppGpp were initially discovered to be synthesized during amino acid starvation, but more recently, it was observed that their synthesis is also induced by protein folding stresses, such as heat or oxidative stress^7^. During heat stress, their accumulation results in increased thermoresistance, inhibition of translation, and a reduction in protein aggregation^7^. (p)ppGpp inhibit translation by binding to specific GTPases (e.g. IF2, RbgA)^15–17^. Furthermore, (p)ppGpp accumulation can lead to decreased transcription of e.g. rRNA and ribosomal protein (r-protein) transcripts by inhibiting GTP biosynthesis and reducing its levels^18,19^. This downregulation of translation is an important component of the response to nutrient scarcity as well as heat shock.

In this study, we aimed to identify new genes or pathways involved in the heat shock response using a genetic screen that reports on fitness. We identified that the Y-complex (RicAFT-complex), which interacts with the endonuclease RNase Y and conveys substrate specificity, plays a role in the heat shock response. We demonstrate further that the Y-complex regulates ribosome levels by initiating ribosome degradation during nutrient depletion or heat shock. Mutants lacking a Y-complex member show increased ribosome levels before and during heat shock, increasing the burden on the PQC system and, in consequence, protein aggregation during heat stress. These findings not only indicate the importance of RNases during stress response but also emphasize that controlling the stability of RNA is an important general mechanism to modify cellular function.

## Results

### The Y-complex is necessary for survival during heat shock

To identify genes that play a role in the heat shock response, we used the BKE single-mutant library^20^, which includes a barcoded deletion cassette for each non-essential gene in *B. subtilis*. All gene deletion mutant strains were pooled, grown at 30°C, and subjected to a mild heat shock at 50°C during the mid-exponential phase, while a control was maintained at 30°C. Relative fitness scores for each gene deletion in the pooled population were calculated by quantifying each barcode at different temperatures (**Fig. 1A, Supplementary Table 1**). The relative fitness of the heat shocked cultures compared to the cultures grown at 30°C was used to identify genes whose deletion results in a change in fitness during heat shock (**Fig. 1A-B, Supplementary Table 1**).

**Figure 1.**
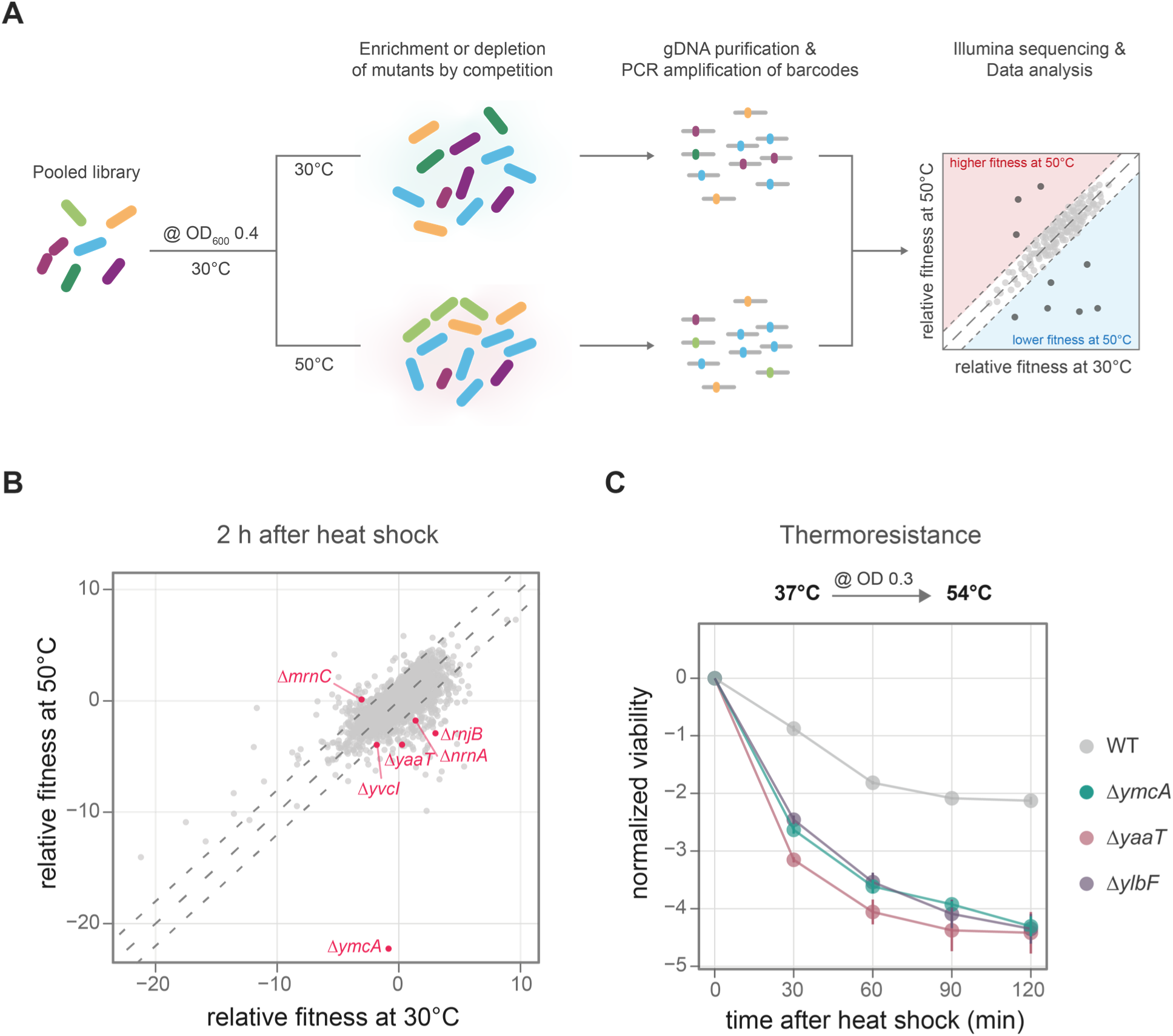
Genetic screening for genes involved in the heat shock response. **A)** Genetic screening experiment: the pooled library of deletion mutants (BKE) was grown at 30°C; when it reached an OD_600nm_ of 0.4, one fraction was exposed to 50°C while the other was maintained at 30°C. Genomic DNA (gDNA) was extracted from the library, and barcodes were amplified and sequenced using Next-Generation Sequencing. The fitness of each mutant at the mentioned temperatures and time points was calculated using BEANcounter. **B**) Relative fitness of each deletion mutant at 50°C (*y*-axis) and 30°C (*x*-axis) two hours post-temperature shift. Each point represents a deletion mutant, with mutants lacking RNase-related genes highlighted in magenta. The dotted lines indicate the range in which the relative fitness between the conditions differed by less than 2. Data represent the average of three biological replicates. A negative relative fitness value means that the abundance of the deletion mutant, compared to the rest of the population, was negatively affected by the treatment, while a positive value means an increased abundance in this condition. **C)** Changes in the viability (log_10_CFU/ml) of WT and *ymcA, ylbF,* and *yaaT* deletion mutants to a severe heat shock at 54°C. Values were normalized to the viability at timepoint 0 min. The data represents the average ± standard error of three biological replicates.

We observed that *B. subtilis* strains lacking the genes coding for the heat shock transcriptional repressors HrcA or CtsR exhibit higher fitness at 50°C compared to 30°C (**Supplementary Fig. 1A**), consistent with their known regulatory role. In contrast, deletion of the genes coding for the DnaK chaperone system, the protease ClpP, and the adaptor protein complex McsA-McsB show greater sensitivity to increased temperatures (**Supplementary Fig. 1B**), demonstrating that our screening approach can identify genes with already known heat-shock phenotypes^21–23^.

The control of transcription initiation by transcription factors has been well established^24^. However, the contribution of RNA stability in stress response is not well understood. Interestingly, six mutants lacking RNases or RNase-regulating genes exhibit differential fitness at 50°C compared to 30°C (**Fig. 1B)**. The deletion of RNase Mini-III (Δ*mrnC)* results in lower fitness at 30°C, whereas its fitness remains unaffected at 50°C. Deletions of RNase J2 (Δ*rnjB)*, the nano-RNase A (Δ*nrnA*), and the RNA pyrophosphohydrolase and (p)ppGpp hydrolase NahA (Δ*yvcI*) exhibit reduced fitness during increased temperatures (**Fig. 1B**).

One of the mutants with the lowest fitness score at 50°C is Δ*ymcA* (**Fig. 1B**). YmcA (RicA) interacts with YaaT (RicT) and YlbF (RicF) to form the Y-complex, which associates with the endonuclease RNase Y for maturation of polycistronic mRNA and riboswitches^25,26^. In addition, we also identified Δ*yaaT* in our screen (**Fig. 1B**). We could confirm the heat sensitivity of all three deletion mutants of the Y-complex genes in single-culture experiments during a severe heat shock at 54°C (**Fig. 1C**).

### Identification of Y-complex-dependent cleavages at late exponential and transition phases

We evaluated the growth of all three strains lacking the Y-complex encoding genes. They exhibit a similar growth defect that begins at the end of the exponential phase and continues during transition phase until the stationary phase (**Fig. 2A**). Given that this complex associates with the endonuclease RNase Y and regulates substrate selection^26^, we investigated Y-complex-dependent cleavages in the late exponential and transition phase (**Fig. 2A**) using the ISCP (Identification of <u>S</u>pecific <u>C</u>leavage <u>P</u>ositions) approach^27^. We compared the coverage of RNA 3’ and 5’ ends in the WT and Δ*ymcA* mutant, and the ends significantly reduced in the Δ*ymcA* were classified as putative Y-complex-dependent cleavages (**Supplementary Table 2**). It is worth noting that deleting any member of the Y-complex results in a significant decrease in the processing of substrate RNA^26^. When a transcript is cleaved, it results in two products: one is quickly degraded by exonucleases, while the other is stabilized^27^. Consequently, it is not always possible to detect both 5’ and 3’ ends after cleavage.

**Figure 2.**
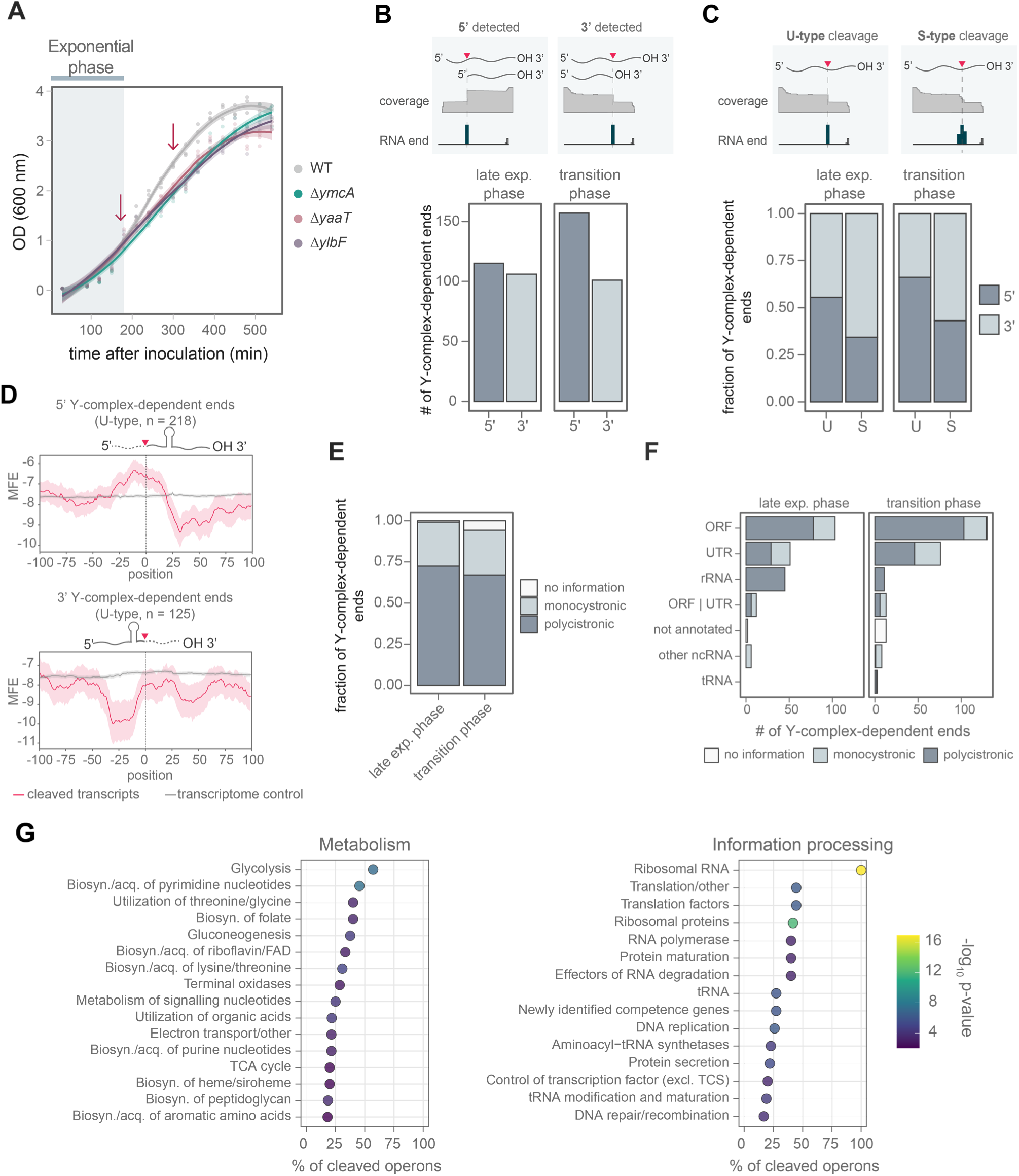
Identification of Y-complex-dependent cleavages at late exponential and transition phase. **A)** Growth curves of WT, *ymcA*, *ylbF*, and *yaaT* deletion mutants in LB at 37°C. The blue bar marks the exponential phase. Arrows indicate sample collection for the ISCP method at the late exponential and transition phases. Three biological replicates are displayed with a smooth curve regression. **B)** Number of RNA 3’ or 5’ ends identified at the two growth phases. **C)** Fraction of identified 3’ or 5’ ends displaying U-type (unique) or S-type (stepped) profiles. **D)** Minimum free energy (MFE) was calculated on 50 nt windows for RNAs cleaved by the Y-complex with U-type ends (in magenta). Transcripts that share the same cleavage motif but are not processed were used as a control (in gray). rRNA genes were excluded from the analysis since they are highly structured. **E)** Fractions of ends identified in genes transcribed in polycistronic or monocistronic RNA. **F)** Count of RNA features where Y-complex-dependent ends are detected at either growth phase. Annotations were obtained from BSGatlas^76^. **G)** Overrepresentation analysis of pathways whose transcripts are cleaved by the Y-complex during the late exponential phase. Operons were assigned to be involved in a pathway if they code for one gene participating in such a pathway. As genes in the same operon can have different functions, transcripts may be linked to multiple roles. Pathway annotations were retrieved from SubtiWiki^79^. The color indicates the statistical significance of the enrichment. Biosyn = biosynthesis, acq = acquisition, excl TCS = excluding two-component systems.

We identified 221 and 258 Y-complex-dependent ends in late exponential and transition phases, with six also detected in a previous study at the mid-exponential phase^26^ (**Supplementary Table 3**). In the late exponential phase, 5’ and 3’ ends are almost equally distributed, yet more 5*’* ends are detected in the transition phase (**Fig. 2B**).

It is known that the Y-complex and RNase Y generate fragments with differential stability in the mid-exponential phase^26^. Here, we evaluated the stability of the resulting fragments in the late exponential and transition phases by calculating the ratio of coverage upstream and downstream of the cleavage site (**Supplementary Table 2**). We observed a trend toward stabilizing downstream fragments. However, upstream fragments can also be stabilized, but to a lesser extent (**Supplementary Fig. 2A**). The ISCP method differentiates processing sites with blunt ends (Unique, U-type) or when it was further processed by exonucleases^27^ (Stepped, S-type). Most S-types were found at the 3’ ends, suggesting further processing by exonucleases, while most U-types were detected at 5’ ends (**Fig. 2C**). This is consistent with the trend to stabilise the downstream fragment (**Supplementary Fig. 2A**).

The Y-complex could target a particular sequence motif, thereby helping to recognize specific substrates. However, a motif search using U-type ends resembles the simple motif described for RNase Y in *B. subtilis*^28^, *S. aureus*^29^, and *S. pyogenes*^27^, being most likely a G upstream of the cleavage site (**Supplementary Fig. 2B**). Given the simple sequence motif, we investigated whether cleavages occur randomly in longer or more abundant transcripts. Notably, approximately 81% of targeted transcripts are cleaved only once, about 16% twice, and very few are cleaved multiple times (**Supplementary Fig. 2C)**. Neither the length nor the abundance of a transcript predicts whether it is cleaved, suggesting that other factors influence where cleavages occur (**Supplementary Fig. 2D-E, Supplementary Table 4**). Recent studies have shown that the cleavage efficiency is multifactorial, depending on the sequence and secondary structures surrounding the cleavage site^28,30^.

As we observed that RNA processing could result in fragments with different stability (**Supplementary Fig. 2A**), we wondered if this could be a consequence of secondary structures protecting them from exonucleases. We used U-type ends and compared transcripts with the same cleavage motif but not cleaved as controls. For the 5’ ends, a strong decrease in the minimum free energy ∼25 nt downstream of the cleavage site indicated a possible secondary structure, while the upstream fragment remained similar to the control (**Fig. 2D**). The upstream fragment’s lower stability may be due to its lack of secondary structure, making it accessible to 3’-5’ exonucleases. For the 3’ ends, the more stable upstream fragment could form a possible secondary structure starting 10 nt upstream of the cleavage site (**Fig. 2D**). The generation of mRNA fragments with different stability might have a regulatory role in gene expression.

### Y-complex-mediated transcript cleavages and mRNA maturation

Most detected Y-complex-dependent ends are inside ORFs, mainly in polycistronic mRNAs (**Fig. 2E-F**), confirming the role of the Y-complex in the maturation and stoichiometry regulation of co-transcribed genes^26^. An example is the *cggR-gapA* transcript, where the Y-complex mediates cleavage in the coding sequence of *cggR* near the stop codon^25,26^. Cleavage renders the *cggR* fragment less stable compared to the *gapA* fragment (**Supplementary Fig. 3A**), thereby uncoupling the transcript levels of these co-transcribed genes. Interestingly, we observed other cleavages generating similar uncoupling of transcripts, such as in the gene encoding the endonuclease YloC (**Supplementary Fig. 3B**, **Supplementary Table 2)**. In addition, cleavages in genes transcribed as monocistronic mRNAs, such as *guaB,* suggest that the Y-complex could initiate their degradation (**Supplementary Fig. 3C**, **Supplementary Table 2**).

We observe Y complex*-*dependent processing in 5’, 3’, and internal UTRs (**Fig. 2F**). Cleavages in 5’ UTR suggest a possible regulatory role by decreasing the abundance of the mRNA containing them, as already described for riboswitches^26,31^, or by producing a shorter version of a 5’ UTR, which could affect gene expression. Processing of 3’ and internal UTR could influence mRNA stability and polycistronic mRNA maturation. We observed that some 3’ UTR cleavages are located upstream of the Rho-independent terminator, decreasing the stability of the upstream fragment and leaving an apparent more stable downstream fragment that includes its terminator structure, as observed for *trxB* (**Supplementary Fig. 3D**). This phenomenon has been observed in *E. coli*, where RNase E cleaves upstream of the Rho-independent terminator^32,33^.

In polycistronic mRNA maturation, we observed that Y-complex processes internal UTRs, decoupling the RNA levels of coding fragments (**Supplementary Fig. 3E-F**). For example, the *bipA*-*ylaH* transcript is cleaved 21 nt after the *bipA* stop codon, rendering the upstream fragment less stable than the *ylaH* fragment (**Supplementary Fig. 3E**). We also observed a cleavage between the *fusA* (EF-G) and *tufA* (EF-Tu) genes, both coding for translation elongation factors (**Supplementary Fig. 3F**).

### The Y-complex targets translation and metabolism-related genes in the late exponential phase

We wondered if the Y-complex primarily processes transcripts bearing genes with specific functions required for switching the growth phase. Therefore, we conducted an overrepresentation analysis and observed that the most significantly enriched categories pertain to “Metabolism” and “Information processing” (**Fig. 2G, Supplementary Fig. 4A, Supplementary Table 5**). Overall, the Y-complex is predominantly involved in the processing of operons encoding genes for carbon metabolism (e.g., glycolysis and gluconeogenesis), nucleotide metabolism (e.g., pyrimidine and purine biosynthesis), and amino acid biosynthesis and utilization (**Fig. 2G**). This allows adjusting the utilization and biosynthesis of nutrients when they become limited. In addition, all transcripts containing rRNA and ∼50% of transcripts carrying genes encoding translation factors and r-proteins are processed by the Y-complex (**Fig. 2G**).

The reduced RNA processing in the *ymcA* deletion strain results in significant changes in the transcriptome (**Supplementary Fig. 4B, Supplementary Table 6-7**). Consequently, transcripts of certain pathways related to metabolism and translation, among others, show maturation events that depend on the Y-complex and also demonstrate differential expression upon deletion of *ymcA* (**Fig. 2G**, **Supplementary Fig. 4B**). This may result from either a direct or indirect effect of decreased RNA maturation events. Our results suggest that the Y-complex and RNase Y are important for transitioning to the stationary phase by processing transcripts related to translation and metabolism, which may impact their stability and/or gene expression.

### The Y-complex targets ribosomes for degradation

Several Y complex-dependent cleavages were detected in the rRNA sequence during both growth phases, most of them being in the 23S rRNA during the late exponential phase (**Fig. 3A**). We detected cleavages in specific helices in the 23S and 16S rRNA, mainly in loops of hairpin structures (**Supplementary Fig. 5A-B**). These helices are accessible and primarily located close to the A-site and the interface between the two ribosomal subunits (**Supplementary Fig. 6**), which could be recognized and cut by the Y-complex with the endonuclease RNase Y. These cleavages will most likely destabilize the ribosome structure. Generating free 5’ and 3’ RNA ends, which are substrates for exonucleases such as PNPase, RNase R, and RNase J, will result in further RNA degradation and ribosome disassembly, releasing the ribosomal proteins as well.

**Figure 3.**
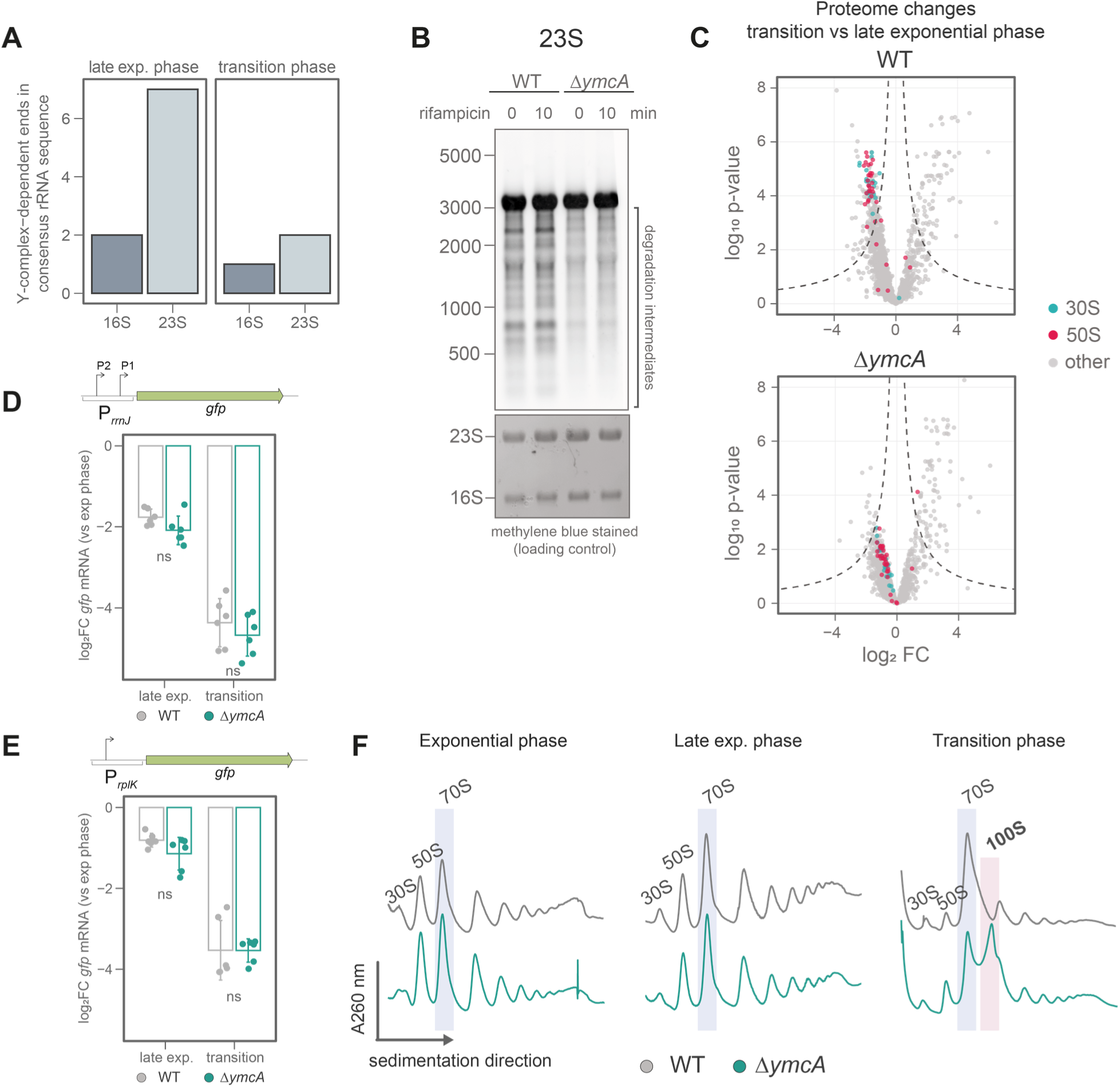
The Y-complex controls ribosome levels by initiating rRNA degradation. **A)** Number of cleavages detected by ISCP in *B. subtilis* rRNA during the late exponential and transition phase. **B)** Northern blot against 23S of WT and Δ*ymcA* cells at the late exponential phase, before and after treatment with 100 µg/ml rifampicin to stop transcription. The methylene blue-stained membrane is shown as a loading control. This image is representative of three biological replicates. **C)** Volcano plot of protein quantification of WT and Δ*ymcA* strains comparing transition to late exponential phase. The proteins composing the small (30S) and major (50S) ribosome subunits are highlighted in cyan and magenta, respectively. The data shows the average of four biological replicates. Activity of the promoter controlling **D)** *rrnJ* (rRNA) or **E)** the *rplK* (r-protein) transcription at late exponential and transition phase compared to the exponential phase in WT and Δ*ymcA*. The promoters were cloned controlling *gfp* transcription, which was measured using RT-qPCR. The *pcp* transcript was used as a reference. Individual values and the average ± standard deviation of three biological replicates and two technical replicates are shown. The statistical significance was tested using student’s *t*-test. ns: *p-*value > 0.05. **F)** Ribosome sedimentation profile in 10-50% sucrose gradients of WT and Δ*ymcA* in exponential, late exponential, and transition phases. The 70S and 100S ribosomes are highlighted with a blue or red box, respectively. The data is representative of three biological replicates.

We confirmed the observed rRNA degradation via northern blots of 23S or 16S rRNA (**Fig. 3B**, **Supplementary Fig. 7A**). Deletion of *ymcA* strongly reduces the presence of rRNA degradation intermediates during the late exponential phase. As ribosomes are assembled co-transcriptionally^34^, we inhibited transcription by adding rifampicin to cultures at the late exponential phase. We observed that stopping transcription does not affect rRNA degradation of the 23S and even raises the levels of 16S degradation intermediates, indicating that mature ribosomes are degraded (**Fig. 3B**, **Supplementary Fig. 7A**). The Y-complex protein YaaT interacts more stably with RNase Y in the membrane than YmcA and YlbF^35^. We tagged YaaT with a C-terminal FLAG and identified interacting proteins during the late exponential phase by pull-down (**Supplementary Fig. 7B, Supplementary Table 8**). We observed that YaaT captures other Y-complex members (YmcA, YlbF), RNase Y, and components of the dynamic degradosome complex, including enolase and PNPase. Interestingly, r-proteins are also observed in the pull-down experiment, suggesting the Y-complex might target ribosomes to RNase Y for initiation of rRNA degradation after exponential growth (**Supplementary Fig. 7B**).

Cleavages within mature rRNA can trigger ribosome disassembly and degradation, which, during the transition phase, coincides with translation downregulation through reduced synthesis and inhibition of ribosome activity. This suggests that the Y-complex may be involved in reducing ribosome levels to accommodate the new, slower growth rate. Therefore, we quantified ribosome content by measuring the abundance of all r-proteins via mass spectrometry (**Supplementary Table 9**). When *B. subtilis* shifts from exponential to the transition phase, we observed a coordinated reduction in r-protein levels (**Fig. 3C**), indicating a decrease in ribosome levels. This reduction is highly impaired in Δ*ymcA* (**Fig. 3C**), where the levels of r-proteins do not decline to the same degree as in the WT, likely due to the reduced degradation of ribosomes.

Accumulation of (p)ppGpp leads to inhibition of transcription of rRNA and r-proteins^19,36^ upon shortages of nutrients during the transition to the stationary phase. To test if the deletion of *ymcA* is interfering with this transcriptional downregulation, we measured the promoter activity of the rRNA promoters *rrnJ* and *rrnB* and r-protein promoters of *rplK*, *rplJ*, and *rpsJ*. All these promoters show similar downregulation in the late exponential and transition phases in both WT and Δ*ymcA* strains (**Fig. 3D-E**, **Supplementary Fig. 7C-E**), confirming that their transcriptional control remains unaffected and is not the cause of the increased ribosome amounts in the Δ*ymcA* strain.

Notably, we observed that the increased levels of ribosomes in a Δ*ymcA* strain led to the accumulation of hibernating 100S ribosomes during the transition phase (**Fig. 3F**). This 100S is no longer observed after the deletion of the gene encoding the hibernation factor *hpf*, which triggers their formation^37,38^ (**Supplementary Fig. 7F**). The presence of hibernating ribosomes at this time point indicates that functional and superabundant ribosomes in the Δ*ymcA* strain are sequestered away from the actively translating pool through hibernation.

### (p)ppGpp and the Y-complex play a crucial role in regulating the levels of ribosomes

Since ribosome levels and translation are lowered by (p)ppGpp under nutrient limitation or other stresses, we wondered if there is a genetic interaction between the two systems. Deleting *ymcA* in a strain that cannot synthesize (p)ppGpp [(p)ppGpp^0^] leads to an even stronger growth defect during the transition phase (**Fig. 4A**). The (p)ppGpp^0^ strain, like Δ*ymcA*, shows impaired reduction of ribosome levels during the transition phase (**Fig. 4B**). The double mutant (p)ppGpp^0^ Δ*ymcA* shows almost no reduction in ribosome quantities during this growth phase (**Fig. 4B-C**). The same is observed when the r-protein mass fraction is calculated at the transition phase (**Fig. 4D**). These experiments suggest that the Y-complex and (p)ppGpp control ribosome levels and translation independently during post-exponential growth. The absence of regulation by both systems results in a severe growth defect.

**Figure 4.**
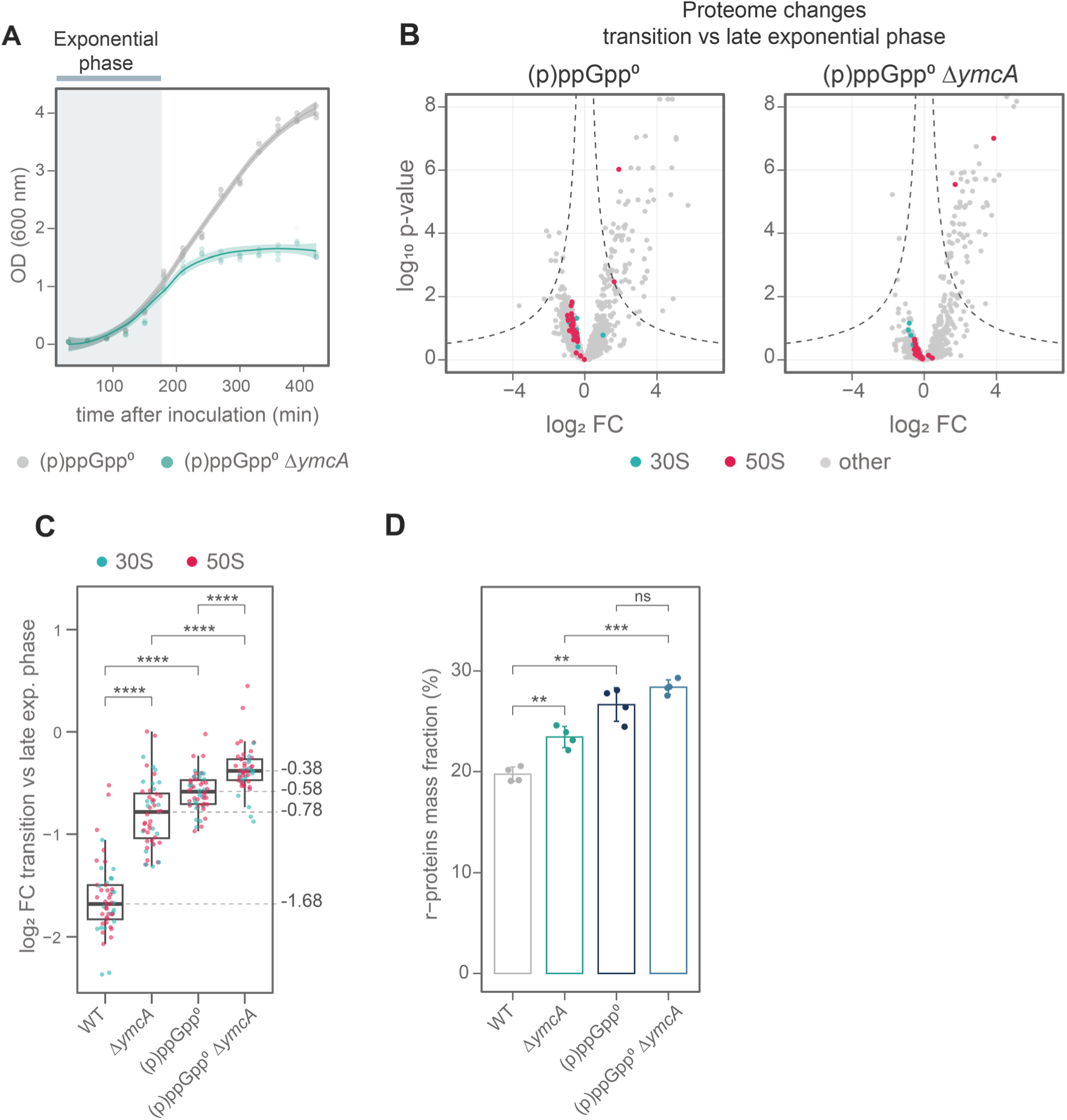
The alarmones (p)ppGpp and the Y-complex are important for controlling ribosome levels. **A)** Growth curve of (p)ppGpp^0^ and (p)ppGpp^0^ Δ*ymcA* strains in LB at 37°C. The blue bar marks the exponential phase. Three biological replicates are displayed with a smooth curve regression. **B)** Volcano plot of proteomic changes of (p)ppGpp^0^ and (p)ppGpp^0^ Δ*ymcA* cultures comparing transition to exponential phase. The proteins composing the small (30S) and major (50S) ribosome subunits are highlighted in cyan and magenta, respectively. The data shows the average of four biological replicates. **C)** Comparison of r-proteins downregulation in WT, Δ*ymcA,* (p)ppGpp^0,^ and (p)ppGpp^0^ Δ*ymcA* during the transition phase compared to the late exponential phase. The boxplot represents the interquartile range (IQR) and the median in the center. Whiskers show the variability outside quartile 1 (Q1) and Q3 and were calculated as Q1-1.5*IQR and Q5+1.5*IQR, respectively. Proteins are highlighted like in **B**. Alternative r-proteins were excluded from the analysis. The statistical significance was tested using paired Wilcoxon test. ****: *p-*value ≤ 0.0001. **D)** Proteome fraction (in %) of r-proteins mass, measured by mass spectrometry. The r-protein mass fraction was calculated for the indicated strains. Individual values and the average ± standard deviation of three biological replicates are shown. The statistical significance was tested using student’s *t*-test. ns: *p-*value > 0.05, **: *p-*value ≤ 0.01, ***: *p-*value ≤ 0.001.

### Increased ribosome levels lead to protein aggregation during heat shock

Since we discovered the Δ*ymcA* strain in a competition experiment during heat shock, we explored whether the Y-complex is involved in lowering ribosome levels upon proteotoxic stress. We evaluated the degradation of 23S and 16S rRNA by the Y-complex before and after heat shock (**Fig. 5A**, **Supplementary Fig. 8A**). After 15 minutes at 50°C, degradation intermediates are observed in the WT but not in Δ*ymcA*, confirming a role of the Y-complex in degrading ribosomes during heat stress (**Fig. 5A**). Comparing r-protein mass fractions before and during heat shock shows that both WT and Δ*ymcA* downregulate ribosome levels during the experiment. Overall, there are increased levels of r-proteins in the Δ*ymcA* strain already before the heat shock (**Fig. 5B**), which suggests a role of the Y-complex in maintaining ribosome levels during normal growth. Notably, the levels and induction of known heat-shock-regulated proteins are not affected in the *ymcA* deletion (**Fig. 5C, Supplementary Table 10, Supplementary Fig. 8B**). This suggests that chaperone and protease induction during heat stress in the Δ*ymcA* strain is intact.

**Figure 5.**
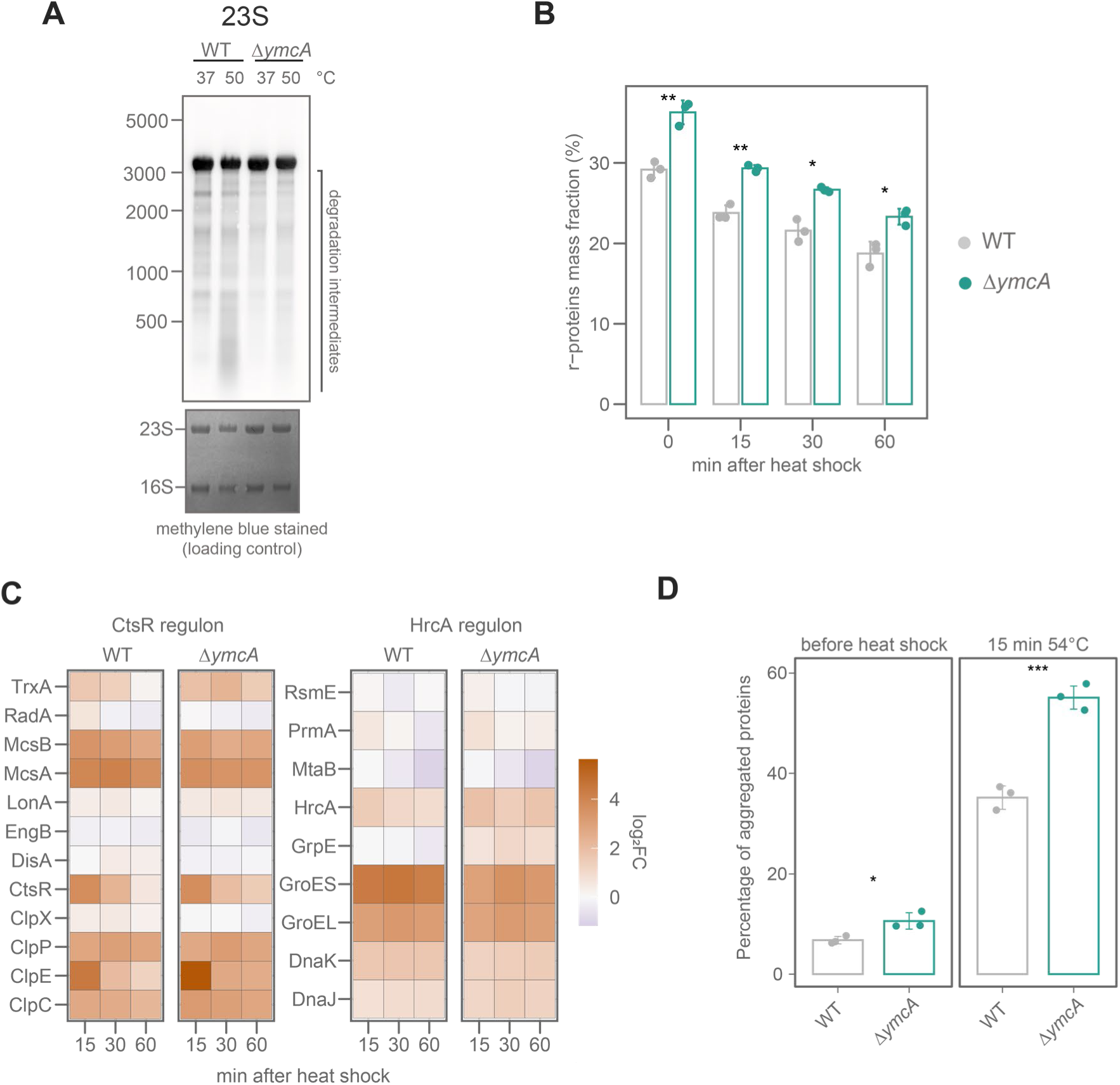
Control of ribosome levels and protein homeostasis in *ymcA* mutant during heat shock. **A)** Northern blot against 23S of WT and Δ*ymcA* cells at exponential phase before or after a heat shock. The methylene blue-stained membrane is shown as a loading control. This image is representative of three biological replicates. **B)** Proteome fraction (in %) of r-proteins mass, measured by mass spectrometry. Individual values and the average ± standard deviation of three biological replicates are shown. The statistical significance was tested using student’s *t*-test. *: *p-*value ≤ 0.05, **: *p-*value ≤ 0.01. **C)** Induction of CtsR and HrcA regulons after heat shock (50°C) in WT and Δ*ymcA*. Proteins were measured by mass spectrometry. The color bar shows the log_2_ fold change compared to the control before heat shock. **D)** Percentage of aggregated proteins before and after a 15 min heat shock at 54°C in WT and Δ*ymcA*. Individual values and the average ± standard deviation of three biological replicates are shown. The statistical difference was tested using student’s *t*-test. *: *p-*value ≤ 0.05, **: *p-*value ≤ 0.01, ***: *p-*value ≤ 0.001.

The downregulation of translation is crucial to limit the increased burden of misfolding and aggregation-prone nascent polypeptide chains on the PQC system. In agreement, a (p)ppGpp^0^ strain shows increased aggregate formation during heat shock^7^. We quantified the relative amount of aggregated proteins before and after 15 minutes of severe heat shock at 54°C and observed that the Δ*ymcA* strain contained significantly higher amounts of aggregated proteins already prior to heat exposure (**Fig. 5D**). This is consistent with the observed increased ribosome levels of the Δ*ymcA* strain already without heat shock (time point 0 in **Fig. 5B**). After 15 minutes of heat shock, Δ*ymcA* shows 20% more aggregated proteins than WT. The (p)ppGpp^0^ Δ*ymcA* mutant exhibits even higher protein aggregation than Δ*ymcA* or (p)ppGpp^0^ strains alone, further supporting that (p)ppGpp and the Y-complex act independently on the regulation of ribosome levels and translation (**Supplementary Fig. 8C**). Mutants of the Y-complex are also strongly affected by other proteotoxic stresses, such as osmotic and oxidative stress, but not by ethanol (**Supplementary Fig. 8D**), which further demonstrates the importance of controlling ribosome levels under proteotoxic stress conditions.

We observed that the Y-complex globally controls the RNase Y-mediated maturation of transcripts and is involved in adjusting ribosome levels to the growth state, allowing the endonuclease RNase Y to initiate the processing and degradation of rRNA and r-protein transcripts. The absence of the Y-complex results in an abundance of idle ribosomes, which interfere with the downregulation of translation and increase sensitivity to protein folding stresses.

## Discussion

Bacterial adaptation to environmental challenges depends on their ability to quickly change gene expression. Beyond the well-known regulation of transcription initiation, we observed here that post-transcriptional mechanisms, which influence RNA stability or translation efficiency, also contribute to the rapid and tailored cellular response. In this context, the control of RNases plays a critical role, since we show that the fitness of *B. subtilis* during heat shock is critically dependent on certain non-essential RNases and the Y-complex, which controls the substrate selection of the quasi-essential endonuclease RNase Y^26^ (**Fig. 1B-C**). Together, our findings demonstrate that the Y-complex – RNase Y system is a regulatory hub that modulates gene expression and the cell’s translational capacity during stress response.

Our RNA-seq analyses reveal that the Y-complex may regulate transcripts involved in a wide range of metabolic pathways, developmental processes, stress responses, and translation (**Fig. 2G, Supplementary Fig. 4A-B, Supplementary Tables 2, 6-7**). RNA processing by the Y-complex-guided RNase Y can create new transcript isoforms^26^ or modify regulatory elements such as UTRs^26,31^(**Fig. 2F**, **Supplementary Fig. 3**), thereby tailoring gene expression to specific growth conditions. This wide regulatory scope of the Y-complex could explain the known pleiotropic phenotypes in these mutants, like decreased sporulation, competence, and defective biofilm formation^39–41^.

Among these diverse functions in RNA processing, we discovered that the Y-complex initiates ribosome degradation at the end of exponential growth by directing RNase Y to cleave 23S and 16S rRNA (**Fig. 3A-B**, **Supplementary Fig. 7A-B**), thereby decreasing the levels of ribosomes. This finding is consistent with previous suggestions that RNase Y can degrade rRNA in aging *B. subtilis* spores^42^.

The exact nature of how the Y-complex (YmcA, YlbF, and YaaT) associates with RNase Y is an active subject of research. Recent evidence indicates that YmcA and YlbF transfer YaaT to RNase Y, forming a stable, membrane-localized YaaT-RNase Y complex^35^. However, the deletion of any of the Y-complex members leads to a strong decrease in the processing of target mRNA^26^, indicating that all Y-complex proteins are equally indispensable for RNA processing. Based on the dynamic localization of YmcA and YlbF between the cell membrane and the cytosol^43^, we propose that these Y-proteins may also recognize substrates, including ribosomes, and direct them to RNase Y for cleavage.

The two [4Fe-4S]^+2^ clusters in the Y-complex^44^ likely act as sensors for cellular growth, energy, and stress, affecting how the complex controls RNase Y. This regulatory complex is localized at the cell membrane, where RNase Y forms a dynamic degradosome-like network with other RNases, helicases, and glycolytic enzymes^45–47^. This spatial co-localization may enable a sequential degradation process: the Y-complex-guided RNase Y performs initial cleavage on targets like rRNA, making fragments susceptible to further degradation by exonucleases such as PNPase, and RNase J as observed with YaaT pull-down experiments (**Supplementary Fig. 7B**). Here, we propose that the Y-complex targets ribosomes to RNase Y and probably other members of the membrane-localized degradosome-like network (**Fig. 6**). This suggests complex spatial control of RNA processing and ribosome decay, separate from transcription and translation in the cytoplasm.

**Figure 6.**
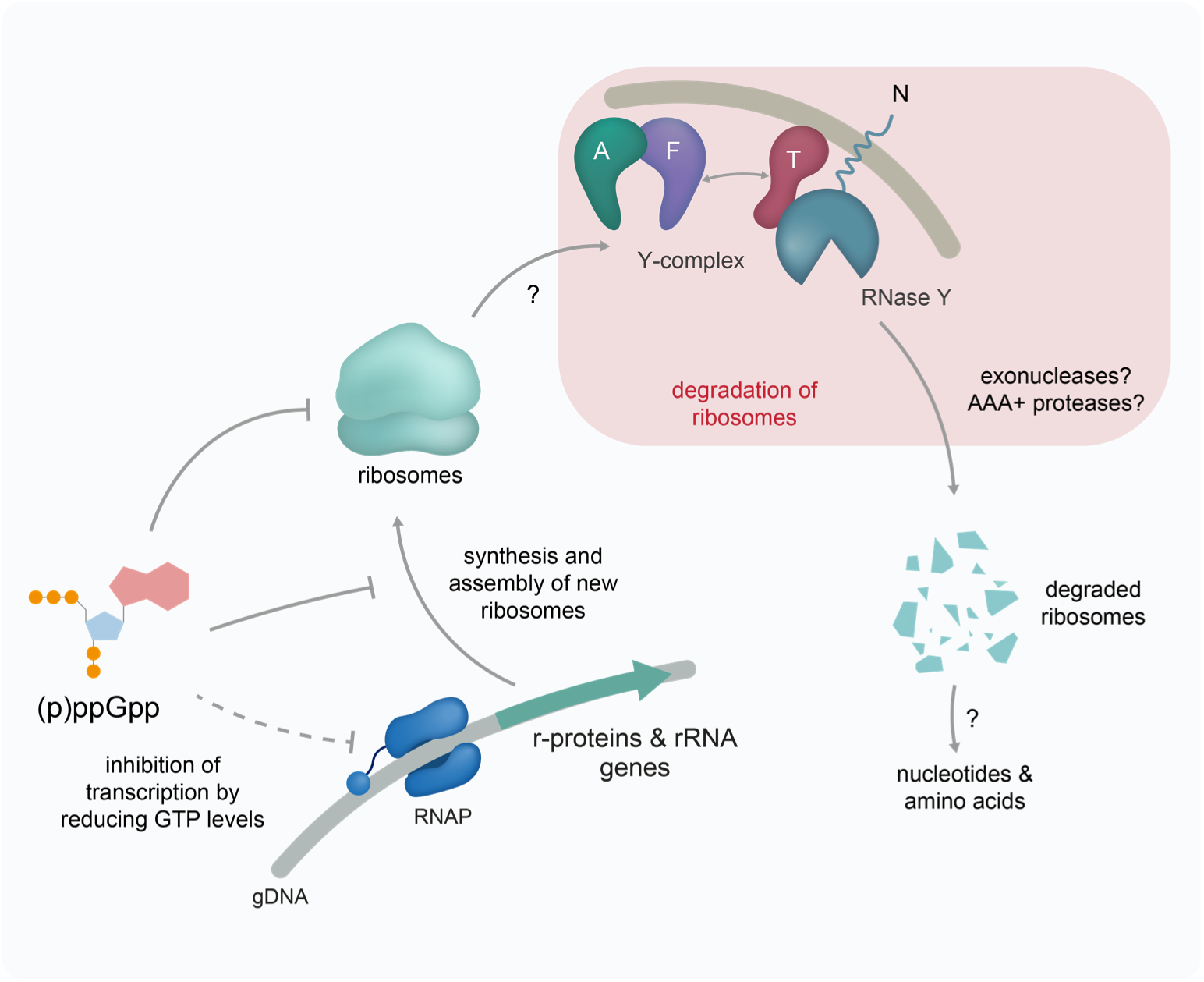
Control of ribosome levels by (p)ppGpp and the Y-complex. The alarmone (p)ppGpp reduces the biogenesis of new ribosomes by indirectly inhibiting the transcription of rRNA and r-proteins mRNA. (p)ppGpp inhibits GTPases that are involved in new ribosome assembly. Additionally, (p)ppGpp reduces translation by binding to specific GTPases. The Y-complex can initiate the degradation of ribosomes, which need to be transported to the membrane by unknown mechanisms. Exonucleases and AAA+ protease complexes might assist in ribosome degradation for the complete degradation of rRNA and r-proteins, which could partially replenish the nucleotide and amino acid pools during starvation.

Most of the cleavages we observed in rRNA were in the 23S (**Fig. 3A**); however, the Y-complex and RNase Y can also initiate 16S degradation in the late exponential phase and during heat shock (**Fig. 3A**, **Supplementary Fig. 7A, Supplementary Fig. 8A**). Recently, it was observed that the exoribonuclease RNase R alone can degrade 16S rRNA^48^, suggesting that endonucleolytic cleavages might not be strictly necessary for 16S degradation. Since RNase R is induced as part of the general stress response^49^, it suggests a possible stress-specific and Y-complex-independent degradation pathway for 16S rRNA.

Complete ribosome decay requires the concurrent degradation of r-proteins (**Fig. 6**). These proteins might be passively released from the ribosome into the cytoplasm during the degradation of the rRNA. Since this would also change their interaction with rRNA, they might misfold and aggregate, and can subsequently be degraded by AAA+ protease complexes. Alternatively, these AAA+ protease complexes could be actively recruited to the site of ribosome degradation to simultaneously extract and degrade the r-proteins along with rRNA. AAA+ protease complexes such as ClpCP with McsB or the membrane-associated FtsH, which functions in protein homeostasis in *B. subtilis*^50^, could be involved in the proteolysis of the r-proteins (**Supplementary Fig. 7B**).

Strikingly, the Y-complex additionally processes transcripts encoding r-proteins and other translation factors (**Fig. 2G**), such as BipA, EF-Tu, and EF-G (**Supplementary Fig. 3E-F**), potentially regulating their translation or leading to their decay. This is supported by the findings that the Y-complex is necessary to process some r-protein transcripts at the mid-logarithmic phase^26^, and RNase Y depletion in this growth phase leads to an increased abundance of transcripts of r-proteins, such as *rplS, rpsO, rpmA, rpsD, rpsB,* and *rplU*^51^.

The control of translation is an important component of many stress response mechanisms. During nutrient starvation, downregulating translation allows the reallocation of resources to biosynthetic pathways^52^. During proteotoxic stresses, it could relieve the overwhelmed PQC system by reducing the synthesis of new proteins that could potentially aggregate^53^. The stress transcription factor Spx and the alarmones (p)ppGpp regulate ribosome levels in *B. subtilis*, and the lack of both impairs growth at 50°C^7^. Our work establishes a mechanistic link between the regulation of ribosome levels through synthesis and decay. The severe growth defect observed in cells lacking both (p)ppGpp and the Y-complex indicates that both of these systems are important for transitioning from exponential to stationary phase. Furthermore, the absence of their combined regulation of translation results in elevated ribosome levels and increased accumulation of protein aggregates. (**Supplementary Fig. 8C**). In times of nutrient limitation, the collective function of (p)ppGpp and the Y-complex in regulating ribosome levels by inhibiting their synthesis and initiating degradation allows resource allocation to other biosynthetic pathways (**Fig. 6**).

These strategies are not exclusive to *B. subtilis*, as different organisms use various mechanisms to degrade ribosomes. These must specifically recognize the ribosome or its subunits to disassemble them and target rRNA and r-proteins for degradation. For example, *E. coli* cells transitioning into the stationary phase also initiate ribosome degradation by the membrane-associated endonuclease RNase E^54,55^, which is functionally equivalent to RNase Y in *B. subtilis*^56^. RNase E forms the scaffold of the membrane-associated degradosome network in *E. coli*^57^. This could allow for the combined activity of endo- and exonucleases, facilitating the complete degradation of the ribosome^56,58^. In addition, it was observed that *E. coli* r-proteins are degraded by the Lon AAA+ protease during starvation^59^. In eukaryotic cells, nutrient starvation also triggers ribosome degradation via the ubiquitin-proteasome system or via specialized autophagy pathways, such as ribophagy^60^. Recent studies have described how stress- or starvation-induced degradation of the 40S ribosome involves the ubiquitylation of proteins S3 and S5. This allows RIOK3 binding, leading to the decay of 18S rRNA by unknown RNases^61–63^.

In summary, we identified the Y-complex as a key regulator of ribosome abundance in *B. subtilis*. The Y-complex regulates the translational activity by triggering ribosome decay and processing transcripts for new ribosomal components. Our findings contribute to the understanding of ribosome degradation by identifying a direct pathway and illustrating how this process is integrated with other stress response mechanisms to control cellular growth and survival. Further studies are required to understand the molecular signals that trigger ribosome degradation and how the decay of rRNA and r-proteins is temporarily and spatially coordinated.

## Materials and methods

### Growth conditions

Unless stated the contrary, strains (**Supplementary Table 10**) were streaked on LB plates (5 g/l yeast extract, 10 g/l tryptone agar, 10 g/l NaCl, 1.5% agar) containing the required antibiotics for selection (1 µg/ml erythromycin + 25 µg/ml lincomycin, 150 µg/ml spectinomycin or 10 µg/ml kanamycin) and incubated overnight at 37°C. A single colony was inoculated in 3 ml of LB medium and grown overnight at 37°C and 180 rpm, then used as the inoculum for subsequent experiments.

### Strain construction

For constructing the strains Δ*ymcA::erm^R^* and Δ*yaaT::erm^R^*, the loci containing the resistance cassette were PCR-amplified from the respective strains of the BKE library^20^ and transformed in *B. subtilis* WT. The deletions of Δ*rel::kan^R^* and Δ*ylbF::erm^R^* were constructed as described in Ref.^20^. For promoter activity reporter strains, the *gfp*(mut2) gene was cloned downstream of the promoters of interest, using the integrative plasmid pDR111 as a backbone.

For transformation, 600 µl of an overnight culture of the receptor strains were diluted in 5 ml competence medium (17.5 g/l K_2_HPO_4_, 7.5 g/l KH_2_PO_4_, 2.5 g/l (NH_4_)_2_SO_4_, 1.25 g/l tri-sodium citrate x2 H_2_O, 0.5% glucose, 7 mM MgSO_4_, 0.1 mg/ml L-tryptophan, 0.02% casamino acids, 0.22 g/ml ammonium iron citrate) and grown at 37°C, 180 rpm, for 3 hours. Subsequently, 5 ml of starvation medium (17.5 g/l K_2_HPO_4_, 7.5 g/l KH_2_PO_4_, 2.5 g/l (NH_4_)_2_SO_4_, 1.25 g/l tri-sodium citrate x2 H_2_O, 0.5% glucose, 7 mM MgSO_4_) were added and grown for 2 more hours. One ml of this culture was incubated with the DNA to be transformed and grown for 2 hours at 37°C and 180 rpm. Positive clones were selected on LB agar containing the required antibiotics. Genetic modifications were verified using PCR and DNA sequencing.

### Genetic screening during heat shock

The BKE library^20^ was replicated on LB agar plates containing 1 µg/ml erythromycin. Liquid cultures were prepared in 200 µl LB medium supplemented with 1 µg/ml erythromycin and grown for 42 hours at 30°C 200 rpm. All mutant strains were pooled and mixed for 15 minutes using a magnetic stirrer. The pooled culture was centrifuged at 5000 rpm for 10 minutes at 18°C. The pellet was resuspended in LB medium to an OD_600_ of 65 and mixed with an equal volume of 50% glycerol. The pooled library was aliquoted and kept at −80°C.

For each replicate, a tube of the pooled library was quickly unfrozen in a water bath at 30°C and used to inoculate 75 ml LB medium in 500 ml flasks to a starting of OD_600_ 0.05. The culture was grown in a water bath at 30°C, 180 rpm, until OD_600_ 0.4. Fifty ml of culture were split into two 250 ml flasks. One flask was transferred to 50°C, 180 rpm for heat shock. The other flask was used as a control sample and was kept at 30°C. One ml samples were collected for gDNA purification at 0 and 2 hours post temperature shift. Samples were pelleted for 2 minutes at 13,000 rpm and washed once with 500 µl PBS. gDNA was purified using the Wizard® Genomic DNA Purification Kit (Promega, A1120) and stored at −20°C until further processing.

The strain-specific barcodes were amplified using indexed primers (**Supplementary Table 7**) with the Collibri Library Amplification Master Mix (ThermoFisher, A38539050) according to the manufacturer’s instructions, with 100 ng of gDNA as input, for 22 cycles. Barcodes were gel-purified, quantified, pooled equimolarly, and sequenced using the Illumina NovaSeq2 single-read method at the Sequencing Core Facility of the Max Planck Institute for Molecular Genetics (Berlin, Germany).

To calculate the fitness score of each deletion mutant from barcode abundance, BEAN-counter v2.6.1^64^ was used with the following settings. A fixed amplicon read structure was used with an index at position 0, the common primer sequence GCAGGCGAGAAAGGAGAG at position 9, and the mutant barcode at position 27 of read 1. All other settings were used with default values. The sample before heat shock served as a reference to which all other time points were compared. All comparisons were performed with three biological replicates.

Fitness data was further analyzed using the R statistical computing language (v4.3.3). For annotation, protein-related information was fetched using the Uniprot REST API (29 February 2024). Average fitness per gene, condition, and time point was calculated as the arithmetic mean of the fitness score from three biological replicates. Differential fitness Δ*F*, e.g. for comparison of treatment *versus* control at a particular time point, was calculated as Δ*F* = *F*_treatment_ - *F*_control_. A two-sided Student’s t-test was performed to evaluate statistical significance for each gene, and the obtained *p*-value was corrected using the Benjamini-Hochberg procedure^65^.

### Growth curves

Twenty-five ml of LB medium were inoculated with an overnight culture to a starting OD_600_ 0.05 in 250 ml Erlenmeyer flasks. Cultures were grown at 37°C 180 rpm in a water bath. OD_600_ was monitored every 30 minutes. The average and standard deviation of three biological replicates were calculated.

### Heat shock survival

Cultures were inoculated in 25 ml LB medium to OD_600_ 0.05 and incubated in a water bath at 37°C 180 rpm. At OD_600_ 0.3 – 0.4, the cultures were exposed to a heat shock at 54°C for 2 hours, and samples for viability measurement were obtained every 30 minutes. Viability was measured by spotting 4 µl of 1:10 serial dilutions in PBS of each sample on LB agar. The plates were incubated overnight at 30°C, and the colony-forming units (CFU) were counted. The CFU/ml was calculated for each sample, and relative viability was determined using samples before heat shock as a reference.

### Isolation of protein aggregates

Twenty-five ml of culture were harvested before and 15 minutes after heat shock at 54°C by centrifugation for 5 minutes at 10,000 rpm. The pellets were washed once with buffer A (100 mM HEPES pH 8.0, 150 mM NaCl) and lysed by bead-beating in buffer B (100 mM HEPES pH 8.0, 150 mM NaCl, cOmplete EDTA-free protease inhibitor) using 0.1 mm glass beads. Afterwards, samples were centrifuged at 1000 x *g* for 2 minutes to obtain the cell extract. The soluble fraction was separated from protein aggregates by centrifugation for 15 minutes at 11,000 x *g* at 4°C. To remove membrane proteins, the aggregates were resuspended in buffer B containing 1% Triton X-100 and incubated for 1 hour at 4°C. Protein aggregates were recovered by centrifugation at 11,000 x *g*, 4°C for 15 minutes and resuspended and incubated as described above. Protein aggregates were washed again with buffer B + 0.5% Triton X-100, and the detergent was removed in a final wash using buffer B. Pellets containing protein aggregates were resuspended in buffer B + 2% SDS by boiling for 5 minutes at 95°C. Protein concentration was measured using the Rapid Gold BCA Protein-Assay-Kit (Pierce™, A53225). The percentage of aggregated protein was calculated to the total protein content in the cell extract.

### Spot test

Overnight cultures were diluted 1:1000 in LB medium without antibiotics and incubated at 37°C 180 rpm for 7-8 hours. The cultures were diluted to OD_600_ 1 in fresh LB medium in a 96-well plate. Seven serial 1:10 dilutions were prepared in 1x PBS. Four µl of each dilution was spotted on LB agar containing different stressors (concentration indicated in the figures). Plates were freshly prepared before use.

### RNA purification

Samples for RNA purification were collected at the indicated time points: exponential phase (OD_600_ ∼ 0.3), late exponential phase (OD_600_ ∼ 1) and transition phase (100 minutes after late exponential phase). Rifampicin was added to a final concentration of 100 µg/ml for 10 minutes.

Samples were harvested by mixing with the same volume of cold 1:1 acetone:ethanol and centrifugated at 9,000 rpm for 5 minutes, 4°C. RNA purification was performed according to Ref.^27^ with slight modifications: The pellet was washed twice in 1 ml TE-sucrose buffer (50 mM Tris-HCl pH 8.0, 10 mM EDTA, 25% sucrose) and centrifugated at 10,000 rpm for 5 minutes, 4°C. Cells were resuspended in 130 µl lysis buffer (20 mM Tris-HCl pH 8.0, 50 mM EDTA, 20% sucrose) supplemented with 3.2 mg/ml lysozyme and incubated on ice for 5 minutes. Two hundred µl of lysis executioner (2% SDS, 1 mg/ml Proteinase K) were added to the samples and incubated at 95°C for 1.5 minutes. Samples were mixed with 1 ml of TRIzol (Ambion) and incubated at room temperature. After 5 minutes, 250 µl chloroform was added to each sample and mixed by vortexing. After 10 minutes at room temperature, the samples were centrifuged for 10 minutes at 11,000 rpm. The RNA was isopropanol-precipitated at −20°C overnight. RNA was pelleted at 11,000 rpm for 20 minutes at 4°C, washed with 1 ml 70% ethanol followed by another centrifugation at 13,000 rpm for 10 minutes at 4°C. The pellet was air-dried and resuspended in 50 µl nuclease-free water.

### ISCP RNA sequencing

RNA sequencing was performed as previously described^66^. In brief, total RNA was treated with TURBO DNase (Ambion, AM2238), depleted of ribosomal RNA using the riboPOOL rRNA depletion kit for *B. subtilis* (siTOOLS; two reactions with 4.5 µg each) and purified using the RNA Clean & Concentrator-5 kit (RCC-5; Zymo Research). RNA 5’ triphosphates were converted to monophosphates using RppH (NEB), and RNA was recovered using phenol:chloroform:isoamyl alcohol extraction followed by ethanol precipitation. RNA ends were repaired using T4 Polynucleotide Kinase (Thermo Scientific), before RNA was purified using the RCC-5 kit and fragmented using an M220 Focused-ultrasonicator (Covaris) as described by the manufacturer but with a treatment time of 100 s (microTUBE AFA Fiber Snap-Cap). After purification using the RCC-5 kit, sequencing libraries were prepared from the recovered RNA using the NEXTflex Small RNA-Seq Kit v3 (Bioo Scientific) according to the manufacturer’s protocol (including step G). Libraries were purified as described in step I of the NEXTflex Rapid Directional qRNA-Seq Kit (Bioo Scientific) using Agencourt AMPure XP Beads (Beckman Coulter; 0.9x bead-to-sample ratio). Sequencing was performed on an Illumina NovaSeq 6000 in paired-end mode with a read length of 100 bp at the Sequencing Core Facility of the Max Planck Institute for Molecular Genetics (Berlin, Germany).

### Processing of RNA sequencing data

FastQC (v0.11.5) was used to assess the quality of the data. Reads were filtered with a minimum quality score of 10 and a length of at least 18 nt, cleaned from adapter sequences using Cutadapt (v3.5)^67^, and mapped against the *B. subtilis* reference genome (NC_000964.3) using STAR (v2.7.10a_alpha_220314)^68^ in ‘random best’ and ‘end to end’ modes. Resulting BAM files were sorted and indexed with Samtools (v1.9)^69^, and PCR duplication artefacts were removed using UMI Tools (v1.1.0)^70^. Gene counts of annotated RefSeq genes were determined with featureCounts (v2.0.3)^71^ and differentially expressed genes (DEG) were calculated with three replicates per condition and time point using DESeq2 (v1.24.0)^72^. *p*-values were corrected for multiple testing using the Benjamini-Hochberg method^65^. To call DEGs, a threshold of adjusted *p*-value < 0.05 and absolute log2 fold change > 1 was used. Genome coverage profiles (total, 5’ and 3’ ends) were generated using a custom script utilizing the HTSeq library (v2.0.2)^73^. We performed the aforementioned analysis steps using a customized pipeline implemented in Snakemake (v7.14.0)^74^.

### Identification of Y complex cleavage sites

The Y-complex*-*dependent cleavage positions were identified following the previously published procedure^66^. In brief, the genome coverage data was prefiltered with a count per million (cpm) value ≥0.05, and only the RNA ends displaying a cpm ≥ 5 were further analyzed. We carried out differential expression analysis of normalized 5′ and 3′ read ends by comparing WT vs. Δ*ymcA* at the late exponential and transition phase. Y-complex-dependent ends were identified using edgeR (v3.28.0)^75^ with absolute log2 fold change ≥ 1 and FDR < 0.05, and kept only if present in both comparisons. These results were further filtered with additional parameters, i.e., the “proportion of ends (cleavage ratio)” and the “ratio of WT and Δ*ymcA* proportion of ends” as was previously described^66^. Moreover, to reduce the potential set of false positive processing sites even further, we applied an additional cut-off on the “proportion of ends” parameter, i.e., we kept only processing sites with a cleavage ratio of the reference strain (WT) ≥ 95%-percentile of the cleavage ratio of the mutant strain.

### Impact of processing sites on transcript and operon expression

To examine the impact of cleavage sites on transcript and operon expression, as well as other genomic features such as untranslated regions (UTRs) and regulatory elements, we calculated transcript per million (TPM) values from variance-stabilized data for each annotated RefSeq gene (transcript level) and for each operon as defined in the BSGatlas (v1.1)^76^ using the DESeq2 package and a custom Python script. Furthermore, all Y-complex-dependent cleavage sites were mapped to various genomic features, including coding regions, UTRs, terminator regions, and riboswitches, as defined in the BSGatlas^76^ (**Supplementary Table 2**).

### Processing site sequence logo

The sequence logos were generated with Logomaker (v0.8)^77^, specifically for 5’ and 3’ end U-type cleavage sites. For this, we extracted 30 nt sequences centered at the cleavage site and calculated the logos with a GC content of 43.5%.

### RNA secondary structure prediction

The RNA structure surrounding Y-complex-dependent cleavage sites was estimated by calculating the MFE (ΔG in kcal/mol) utilizing RNAfold (v2.6.4)^78^. This analysis employed a sliding window approach with 50 nt sequences, focusing on a 200 nt region centered around the position of interest. The average MFE at each nucleotide was then calculated. To calculate and visualize the background folding energy, we randomly draw potential processing sites from the genome determined with the corresponding 5’ or 3’ end motif and calculate the average MFE at each nucleotide as described above. Potential processing sites were drawn 50 times independently.

### Overrepresentation analysis

Annotations of operons were obtained from BSGatlas^76^ and gene categories from Subtiwiki (24 July 2023)^79^. The overrepresentation analysis was performed using the R statistical computing language (v.4.2.2) and the Bioconductor package Category (v.2.64.0).

### Northern blot

Biotinylated RNA probes were prepared using the HighYield T7 Biotin11 RNA Labeling Kit (UTP-based) (Jena Bioscience, RNT-101-BIOX) from PCR amplified fragments containing the promoter for the T7 RNA polymerase (**Supplementary Table 11**). Probes were purified using the Monarch® Spin RNA Cleanup Kit (NEB, T2040) according to the manufacturer’s instructions. Probes were designed taking into consideration the predicted cleavage sites.

One and a half µg of total RNA was separated on denaturing agarose gels (1% agarose, 20 mM MOPS, 5 mM NaOAc, 1 mM EDTA, 6.6% formaldehyde, pH 7.0) in 1x AGE-NB Running Buffer (20 mM MOPS, 5 mM NaOAc, 1 mM EDTA, 0.74% formaldehyde, pH 7.0). The gel was washed 3 times with milli-Q water and once with 10x SSC buffer, and the nitrocellulose membrane was washed once with milli-Q water and once with 10x SSC. The RNA was transferred using a vacuum blotter at 300 mbar covered with 10x SSC for 1.5 hours. After transfer, the membrane was washed for 2 minutes with 6x SSC, once with water, and dried and crosslinked at 254 nm 0.12 J twice. The membrane was developed with methylene blue, and the staining was removed by washing with 0.2x SSC containing 1% SDS, followed by washing with milli-Q water. The hybridization of 16S or 23S rRNA probes was performed using the North2South Chemiluminescent Hybridization and Detection Kit (Thermo) and North2South Hybridization Stringency Wash Buffer (2X) according to the manufacturer’s instructions. Blots were developed in a Vilber imaging system.

### RT-qPCR

Twenty µg of total RNA were treated with TURBO DNase (Ambion, AM2238) and purified using the Monarch® Spin RNA Cleanup Kit (NEB, T2040). Subsequently, cDNA was prepared using the LunaScript RT Mastermix (NEB, E3025) and 2.5 µg of DNase-treated RNA as input. Quantitative PCR was performed using the Luna Universal qPCR Master Mix (NEB, M3003) in a QuantStudio 5 Real-Time PCR System (Applied Biosystems). To calculate changes in *gfp* transcription, samples from the late exponential and transition phases were compared to those from the exponential phase, with *pcp* serving as a reference gene using the Pfaffl method^80^. Primers used for RT-qPCR are listed in **Supplementary Table 11**.

### Ribosome sedimentation in sucrose gradients

Cultures of WT and Δ*ymcA* were grown until the exponential phase (OD_600_ ∼ 0.3), late exponential (OD_600_ ∼ 1), and transition phase (100 minutes after the late exponential phase). Chloramphenicol was added at a concentration of 100 µg/ml 2 minutes prior to sample collection. Fifty ml of culture were pelleted at 4,500 rpm for 10 minutes at room temperature. Pellets were transferred to ice and resuspended in 2 ml of lysis buffer (20 mM Tris-HCl pH 8.0, 100 mM NH_4_Cl, 10 mM MgCl_2_, 0.4% Triton X-100, 0.4% NP-40, 1 mM chloramphenicol, 100 U/µl DNase I), and flash-frozen. Cells were lysed under cryogenic conditions using 5 cycles of mix-milling at 30 Hz for 1 minute. Lysates (240 µg RNA) were laid over a 10-50% sucrose gradient in 20 mM Tris-HCl pH 8.0, 100 mM NH_4_Cl, 10 mM MgCl_2_, 0.5 mM DTT and resolved at 35,000 rpm for 3 hours at 4°C. Gradients were analyzed using a piston fractionator (Biocomp instruments), and absorbance at 260 and 280 nm was monitored.

### Pulldown of YaaT

Cultures of in-locus C-terminal FLAG-tagged *yaaT* strain were grown until the late exponential phase (OD600 ∼ 1). Two hundred ml of culture were pelleted at 15,000 rpm for 2 min. Pellets were transferred to ice and resuspended in 1 ml of resuspension buffer (20 mM HEPES-KOH pH 7.5, 30 mM NH_4_Cl, 6 mM Mg-Acetate, 0.5 mM TCEP) supplemented with cOmplete EDTA-free protease inhibitor), and flash-frozen. Cells were lysed under cryogenic conditions using 3 cycles of mix-milling at 30 Hz for 1 minute. Lysates were stored at −80°C until further processing. Lysates were quickly melted at 30°C, incubated for 10 min on ice with RNase-free DNase I (Roche), and clarified by centrifugation at 12,000 rpm for 5 min at 4°C. DDM (n-dodecyl β-D-maltoside) was added to a 200 µM final concentration. The lysate (input sample) was incubated with 200 µL of anti-FLAG M2 affinity gel (A2220, Sigma) for 2 h at 4°C with constant shaking. Beads were washed 5 times with resuspension buffer supplemented with 200 µM DDM. Proteins were eluted for 45 min in resuspension buffer supplemented with 200 µM DDM and 150 µg/ml of 3x FLAG peptide (Sigma) at 4°C with constant shaking. Proteins were identified and quantified using DIA-MS.

### Sample preparation for MS-based proteomics

Samples were collected at the indicated time points: late exponential phase (OD_600_ ∼ 1) and transition phase (100 minutes after the late exponential phase). For the proteomic changes during heat shock, cultures were grown at 37°C until OD_600_ ∼ 0.3 and shifted to 54°C. Samples were collected at 0, 15, 30, and 60 minutes after heat shock. Ten ml of culture were collected by centrifugation at 4,500 rpm for 10 minutes, and the pellet was stored at −80°C until further processing.

Cell pellets were resuspended in 1 ml of cold PBS and centrifuged at 13,000 rpm for 5 minutes at 4°C. The pellet was washed once more with PBS, resuspended in 150 µL of lysis buffer (100 mM HEPES pH 8.0, 2% SDS, 20 mM TCEP, 60 mM CAA), and lysed using bead beating. Cell debris was discarded by centrifuging at 13,000 rpm for 5 minutes at 4°C. The supernatant was boiled at 95°C for 5 minutes, cooled down to room temperature, and the protein concentration was estimated using the Rapid Gold BCA protein assay kit (Pierce). Samples were flash-frozen in liquid nitrogen and stored at −80°C until further processing.

All samples were subjected to SP3 sample preparation^81^ on an Agilent BRAVO liquid handling robot. Ten µg of a 1:1 mixture of hydrophilic and hydrophobic carboxyl-coated paramagnetic beads (SeraMag, #24152105050250 and #44152105050250, GE Healtcare) were added for each µg of protein. Protein binding was induced by the addition of acetonitrile to a final concentration of 50% (v/v). Samples were incubated at room temperature for 10 minutes. The tubes were placed on a magnetic rack, and beads were allowed to settle for 3 minutes. The supernatant was discarded, and beads were rinsed three times with 200 µL of 80% ethanol without removing the tubes from the rack. Beads were resuspended in digestion buffer containing 50 mM triethylammonium bicarbonate and both Trypsin (Serva) and Lys-C (Wako) in a 1:50 enzyme-to-protein ratio. Protein digestion was performed for 14 hours at 37°C in a PCR cycler. Afterward, the supernatant was recovered and dried down in a vacuum concentrator.

### Proteomics using Mass spectrometry

All samples obtained during the late exponential and transition phase were analyzed on an Orbitrap Exploris (Thermo Scientific) equipped with a FAIMS Pro device and coupled to a 3000 RSLC nano UPLC (Thermo Scientific). Samples were loaded on a PepMap trap cartridge (300 µm i.d. x 5 mm, C18, Thermo) with 2% acetonitrile and 0.1% TFA at a flow rate of 20 µl/min. Peptides were separated over a 50 cm analytical column (Picofrit, 360 µm O.D., 75 µm I.D., 10 µm tip opening, non-coated, New Objective) that was packed in-house with Poroshell 120 EC-C18, 2.7 µm (Agilent). Solvent A consists of 0.1% formic acid in water. Elution was carried out at a constant flow rate of 300 nl/min using a 60-minute method: 8-30% solvent B (0.1% formic acid in 80% acetonitrile) within 23 minutes, 30-45% solvent B within 3 minutes, 45-98% buffer B within 0.5 minutes, followed by column washing and equilibration. Data acquisition on the Orbitrap Exploris was carried out using a data-dependent acquisition (**DDA**) method in positive ion mode. MS survey scans were acquired from 375-1500 m/z in profile mode at a resolution of 60,000. AGC target was set to 200% at a maximum injection time of 25 ms. Peptides with charge states 2-6 were isolated within a window of 1.2 m/z and subjected to HCD fragmentation at a normalized collision energy of 27%. The MS2 AGC target was set to 200%, allowing a maximum injection time of 22 ms. Product ions were detected in the Orbitrap at a resolution of 15,000. Precursors were dynamically excluded for 20 s. The cycle time was set to 1 s for each of the two FAIMS voltages of −45 and −65 V.

LC-MS/MS analysis of the heat shock samples was performed on an Ultimate 3000 UPLC coupled to a Lumos hybrid mass spectrometer (Thermo Scientific). Peptides were loaded onto a PepMap trap cartridge (300 µm i.d. x 5 mm, C18, Thermo) with 2% acetonitrile and 0.1% TFA at a 20 µl/min flow rate. Peptides were separated over a 50 cm analytical column (Picofrit, 360 µm O.D., 75 µm I.D., 10 µm tip opening, non-coated, New Objective) that was packed in-house with Poroshell 120 EC-C18, 2.7 µm (Agilent). Solvent A consists of 0.1% formic acid in water. Elution was carried out at a constant flow rate of 250 nl/min using a 60-minute method: 8-30% solvent B (0.1% formic acid in 80% acetonitrile) within 23 minutes, 30-45% solvent B within 3 minutes, 45-98% buffer B within 0.5 minutes, followed by column washing and equilibration.

The mass spectrometer was operated in data-independent acquisition (**DIA**) and positive ionization mode. MS1 full scans (375–1500 m/z) were acquired with a resolution of 60,000, a normalized automatic gain control target value of 120%, and a maximum injection time set to 55 ms. Peptide fragmentation was performed using higher energy collision-induced dissociation and a normalized collision energy of 30%. MS2 spectra were acquired with a resolution of 30,000 (first mass 100 m/z) with 8 m/z DIA windows between 400 and 1200 m/z. A normalized automatic gain control target value of 100%, and a maximum injection time of 54 ms.

For pulldown samples, proteins were processed following the SP3 protocol as described above. Label-free DIA analyses of peptides were acquired over 120 min by an Orbitrap Exploris 480 (Thermo Scientific) coupled to a 3000 RSLC nano UPLC (Thermo Scientific). Samples were loaded on a pepmap trap cartridge (300 µm i.d. x 5 mm, C18, Thermo) with 2% acetonitrile, 0.1% TFA at a flow rate of 20 µL/min. Peptides were separated over a 25 cm analytical column (PepSep C18, 75 µm I.D., 1.5 µm). Solvent A consists of 0.1% formic acid in water. Elution was carried out at a constant flow rate of 250 nL/min within 120 minutes. Initially, a two-step linear gradient was applied: 3-30% solvent B (0.1% formic acid in 80% acetonitrile) within 70.5 minutes, 30-45% solvent B within 13 minutes, followed by column washing and equilibration. The column was kept at a constant temperature of 50°C.

The MS was operated in DIA mode for single-injection quantitative measurements of individual samples with the following settings: 60k MS1 resolution, MS1 scan range 375-1600 m/z, 15k MS2 resolution, MS2 scan range 120-1600 m/z, Normalized AGC target of 1000%, maximum injection time 54 ms, and fixed normalized collision energy of 30. 12 m/z precursor isolation windows with optimized window placements from 400.4319 to 1210.8024 m/z.

### Proteomics data analysis

For samples at the late exponential and transition phase, Raw files were processed with MSFragger v3.5 using the default settings. Briefly, peak lists were extracted from raw files and searched using a Uniprot *Bacillus subtilis* database (v220530, taxonomy ID 224308) and a database containing sequences of common contaminants and decoys. Trypsin/P was set as enzyme specificity, allowing a maximum of two missed cleavages. Peptide matches were filtered to 1% FDR using PeptideProphet and Philosopher (v4.4.0).

For samples during heat shock, DIA-NN (DIA-NN v1.8.1) was employed for DIA data analysis. The main DIA-NN report, “report.txt”, was used for all calculations. Fasta file (Bacillus_subtilis_Strain168_UP000001570.fasta) was digested with a maximum of one missed cleavage. Peptide length was restricted from 7 to 30 peptides, and the precursor m/z range was set from 300 to 1800. N-terminal methionine excision was enabled. Cysteine carbamidomethylation was selected as a fixed modification, and methionine oxidation and N-terminal acetylation as variable modifications. The maximum number of variable modifications was set to two. All other parameters were at default settings, including the 1% precursor FDR and enabled match between runs (MBR). The main DIA-NN report, “report.pg_matrix.tsv,” was used for all calculations.

For pulldown samples, raw data analysis was performed using Spectronaut® (Biognosys AG, Zurich, Switzerland) version 19.7.250203.62635 in directDIA+ deep mode using a UniProt *B. subtilis* database (v220530, taxonomy ID 224308). Methionine oxidation and Acetyl (Protein N-term) were set as a variable, and carbamidomethylation on cysteine residues was used as a static modification. The FDR for PSM-, peptide-, and protein-level was set to 0.01. All tolerances were set to dynamic for pulsar searches.

The R statistical computing language was used for data processing (v4.4.1). The main comparison of WT, Δ*ymcA*, (p)ppGpp^0,^ and (p)ppGpp^0^ Δ*ymcA* mutants was based on samples from data-dependent acquisition (DDA) experiments. Raw peptide intensity values from label-free quantitation were imported as tabular output from MSFragger. The MSstats package (v4.12.1)^82^ was used to convert raw intensities to an MSstats experiment. In this process, peptide quantification data was log_2_-transformed, median-normalized, and summarized to protein abundance. All features mapping to a protein were used for quantification, Tukey’s median polish was used as a summarization strategy, and missing values were imputed using MSstats’ default strategy. Feature abundance and inferred protein quantity were visually inspected using profile plots. Experimental groups were compared using MSstats’ *groupComparison* function, and the similarity of conditions and replicates was inspected using principal component analysis (PCA, function *prcomp*). For annotation, protein-related information was fetched from Uniprot using the Uniprot REST API (14 June 2024). In addition to the DDA experiments, a time series of WT and Δ*ymcA* mutants was obtained using data-independent acquisition (DIA) proteomics. To quantify relative changes, the log_2_ fold change of each time point was calculated using the zero time point as a reference. To quantify absolute protein abundance for the ribosomal proteome sector, the coarse-graining strategy was used as in Ref.^83,84^. Briefly, two protein sectors (groups) were created containing all annotated proteins of ribosomal subunits. Raw MS1 intensities for all proteins of the respective group and time points were summed up and divided by the total sum of protein intensities, yielding the protein mass fraction per sector.

## Data availability

Amplicon-seq and RNA-seq raw reads have been deposited at the European Nucleotide Archive (ENA) under accession number PRJEB90661. Raw proteomic data have been deposited at the PRIDE database under the accession number PXD065465 for DDA proteomics of late exponential and transition phase samples. The DIA proteomics raw data from the heat shock experiment can be accessed using the accession number PXD065462.

## Author contributions

K.T., K.D., and F.A.C.; Conceptualization. K.D., F.A.C., K.T., R.A.-B., T.F.W., and K.A.: Methodology. F.A.C., K.D., K.S., V.S., K.H., and F.K.: Investigation. K.D., F.A.C., R.A.-B., M.J., K.A., S.R., and K.T.: Formal analysis. F.A.C., K.D., R.A.-B., M.J.: Visualization. F.A.C., K.D., and K.T.: Writing – Original Draft. K.T. and E.C.: Funding acquisition. All authors: Writing – Review & Editing.

## Acknowledgments

We thank Gert Bange (Philipps-Universität Marburg) and Rainer Nikolay (Max Planck Institute for Molecular Genetics) for critically reading the manuscript and providing valuable feedback. This work was funded by the Deutsche Forschungsgemeinschaft (Tu106/8, SPP1879 for K.D. and K.T., Leibniz Prize for E.C.) and the Max Planck Society (for E.C. and K.T.).

---

**Supplementary Figure 1.**
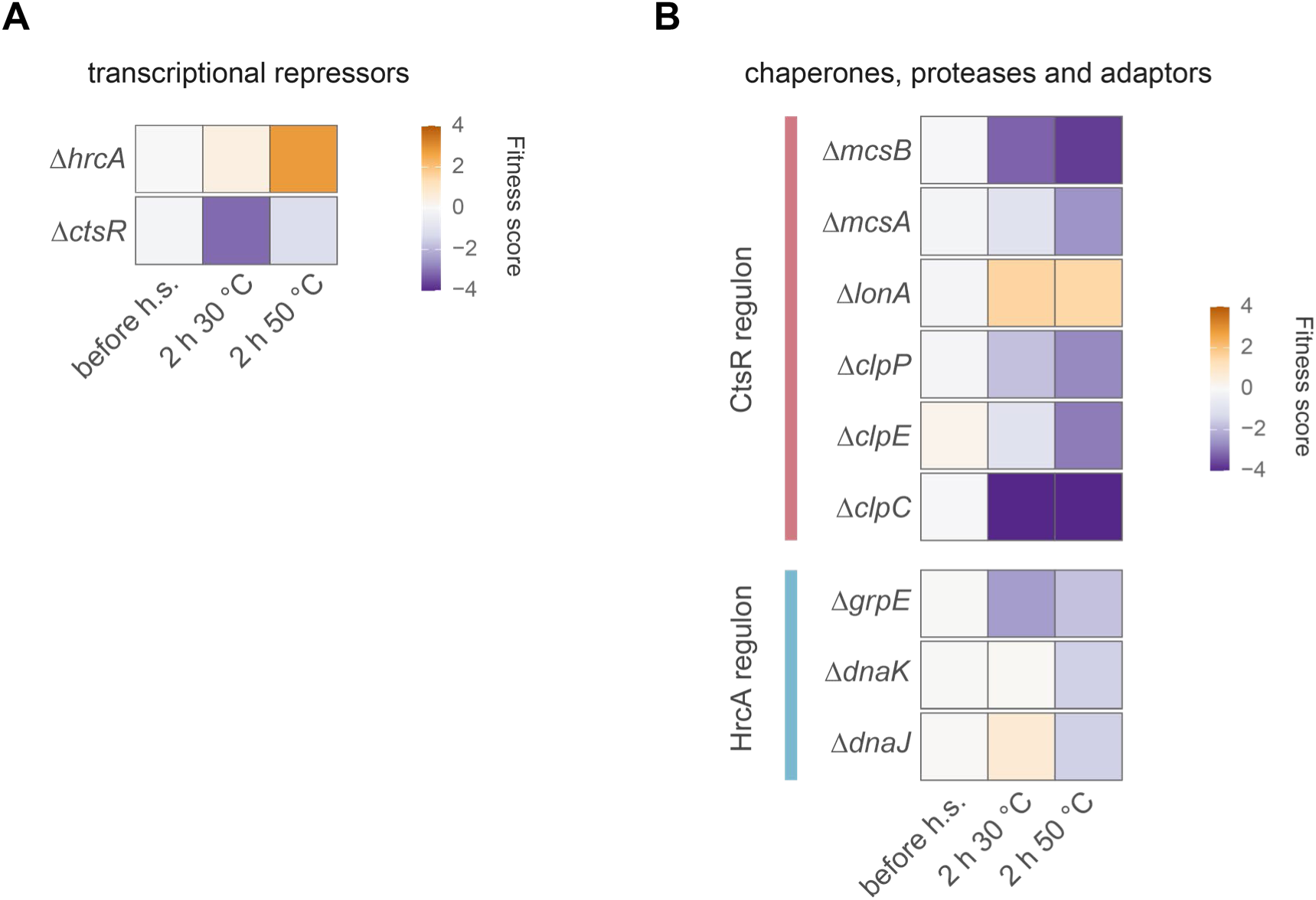
Fitness of selected deletion mutants during heat shock in the genetic screening. Heatmap of the relative fitness of **A)** heat shock transcriptional repressors, and **B)** chaperones, proteases and adaptors. The color represents the average relative fitness of three biological replicates. A negative relative fitness value means that the abundance of the deletion mutant, compared to the rest of the population, was negatively affected by the treatment, while a positive value means an increased abundance on those conditions. h.s. = heat shock.

**Supplementary Figure 2.**
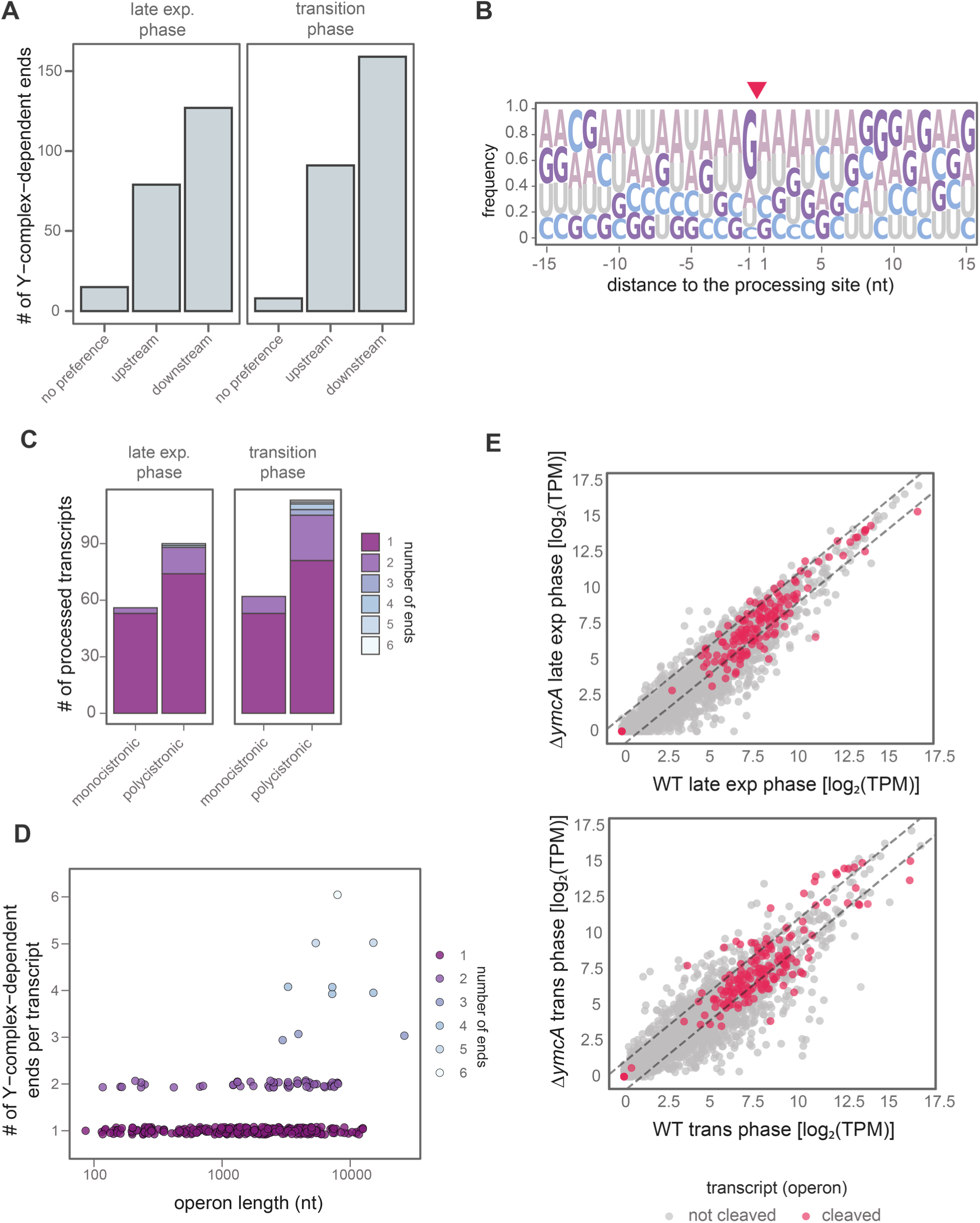
Recognition motif, length, or expression levels do not determine the Y-complex substrates. **A)** Stabilization of the resulting fragments after Y-complex processing, the coverage ratio in a window of 5 nt before and after the processing site was calculated and used as a proxy of the stabilization of the resulting fragments. The predicted stability correlates with the end detected in the ISCP method. **B)** Motif search using 5’ and U-type ends. The arrow indicates the cleavage site. **C)** Number of Y-complex-dependent cleavages detected per monocistronic or polycistronic transcripts in the late exponential or transition phase. **D)** Number of cleavages detected by operon length of the transcript. The color indicates the numbers of Y-complex-dependent ends detected per transcript **E)** RNA levels of WT and Δ*ymcA* mutant transcripts at late exponential and transition phases. Transcripts processed by the Y-complex are highlighted in magenta. Transcript annotations were obtained from BSGatlas.

**Supplementary Figure 3.**
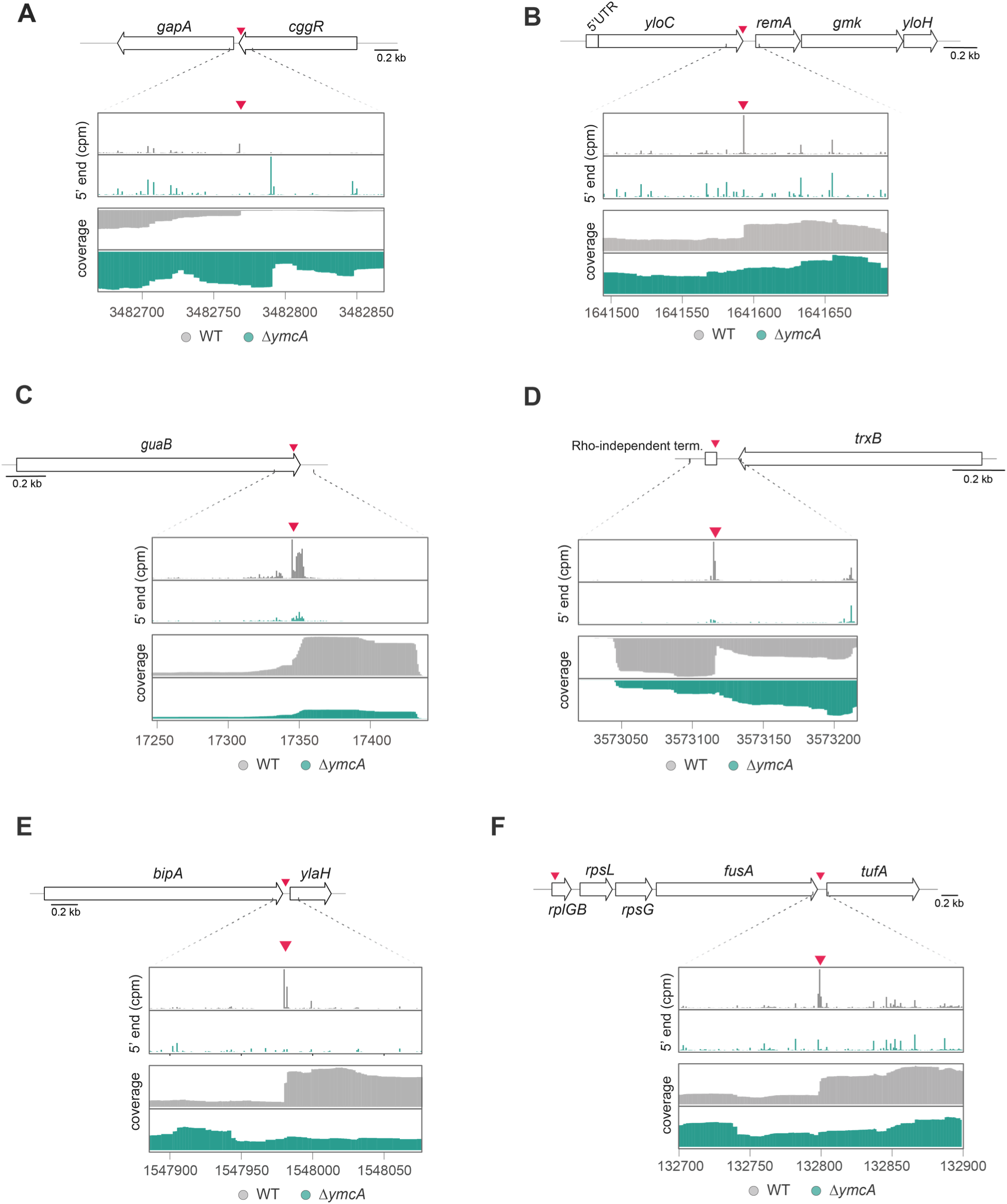
The Y-complex processes mRNA in UTRs and coding regions. RNA end counts per million (cpm), and RNA-seq coverage of selected transcripts in WT and Δ*ymcA.* The arrow indicates the Y-complex-dependent processing site. **A)** The *gapA-cggR* transcript was used as a positive control. **B)** Processing in the coding region of *yloC* in the polycistronic mRNA *yloC-remA-gmk-yloH*. **C)** Processing in the coding region of the monocistronic transcript of *guaB*. **D)** Processing at the 3’ UTR of *trxB,* near the Rho-independent terminator. Processing in the internal UTR of translation-associated transcripts like **E)** the *bipA-ylaH* transcript, and **F)** between *fusA-tufA*.

**Supplementary Figure 4.**
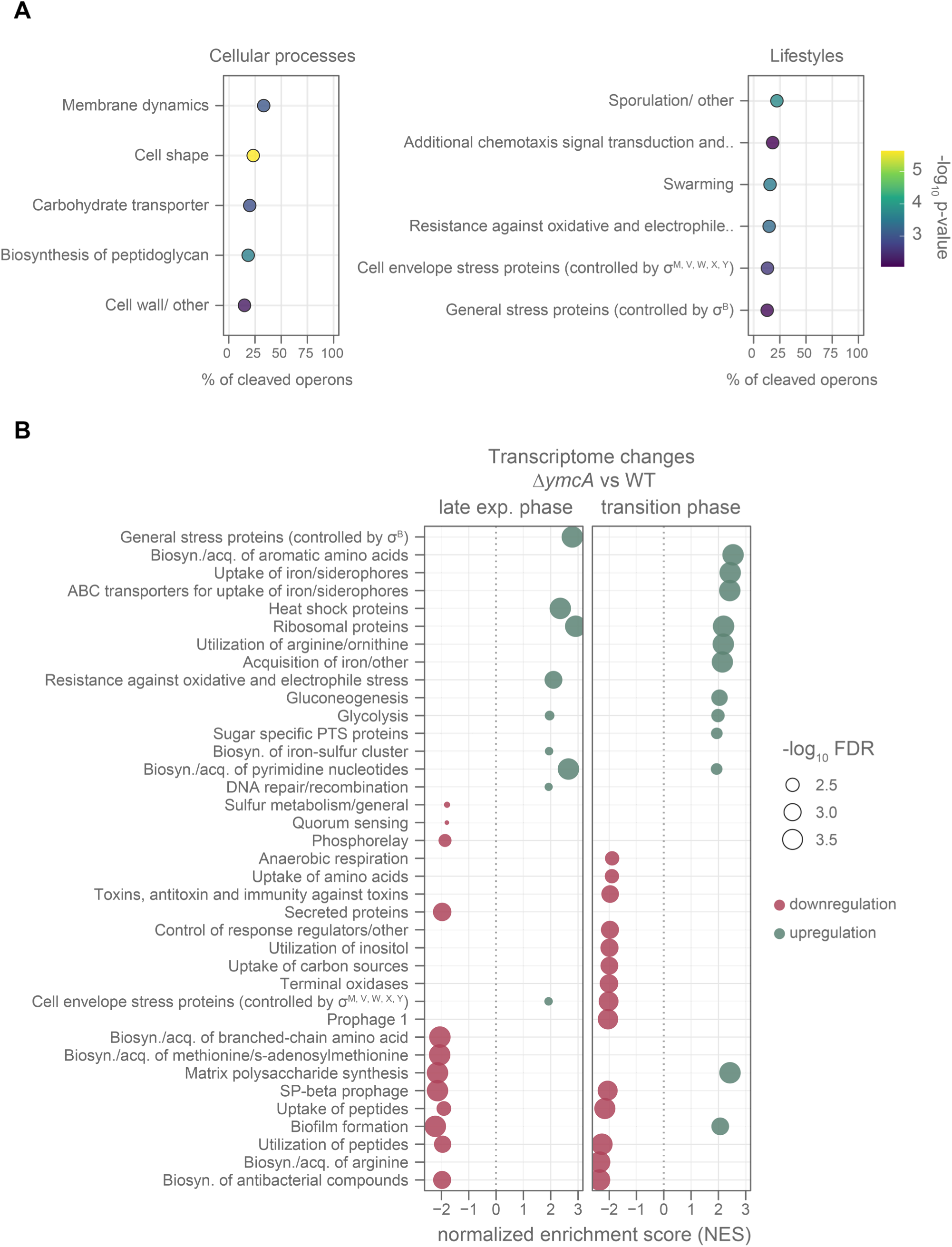
mRNA processing by the Y-complex and effect on the transcriptome. **A)** Overrepresentation analysis of pathways whose transcripts are cleaved by the Y-complex during the late exponential phase. Operons were assigned to be involved in a pathway if they code for one gene participating in such a pathway. Pathway annotations were retrieved from SubtiWiki^79^. The color indicates the statistical significance of the enrichment. **B)** Gene set enrichment analysis (GSEA) of differentially expressed genes at the late exponential and transition phase when comparing Δ*ymcA* to WT. The size of the dot represents the statistical significance. The normalized enrichment score is displayed on the *x*-axis; positive values mean upregulation; meanwhile negative values mean downregulation. GSEA results were prefiltered for pathways showing and enrichment with an FDR ≤ 0.01. Biosyn. = Biosynthesis, acq. = Acquisition.

**Supplementary Figure 5.**
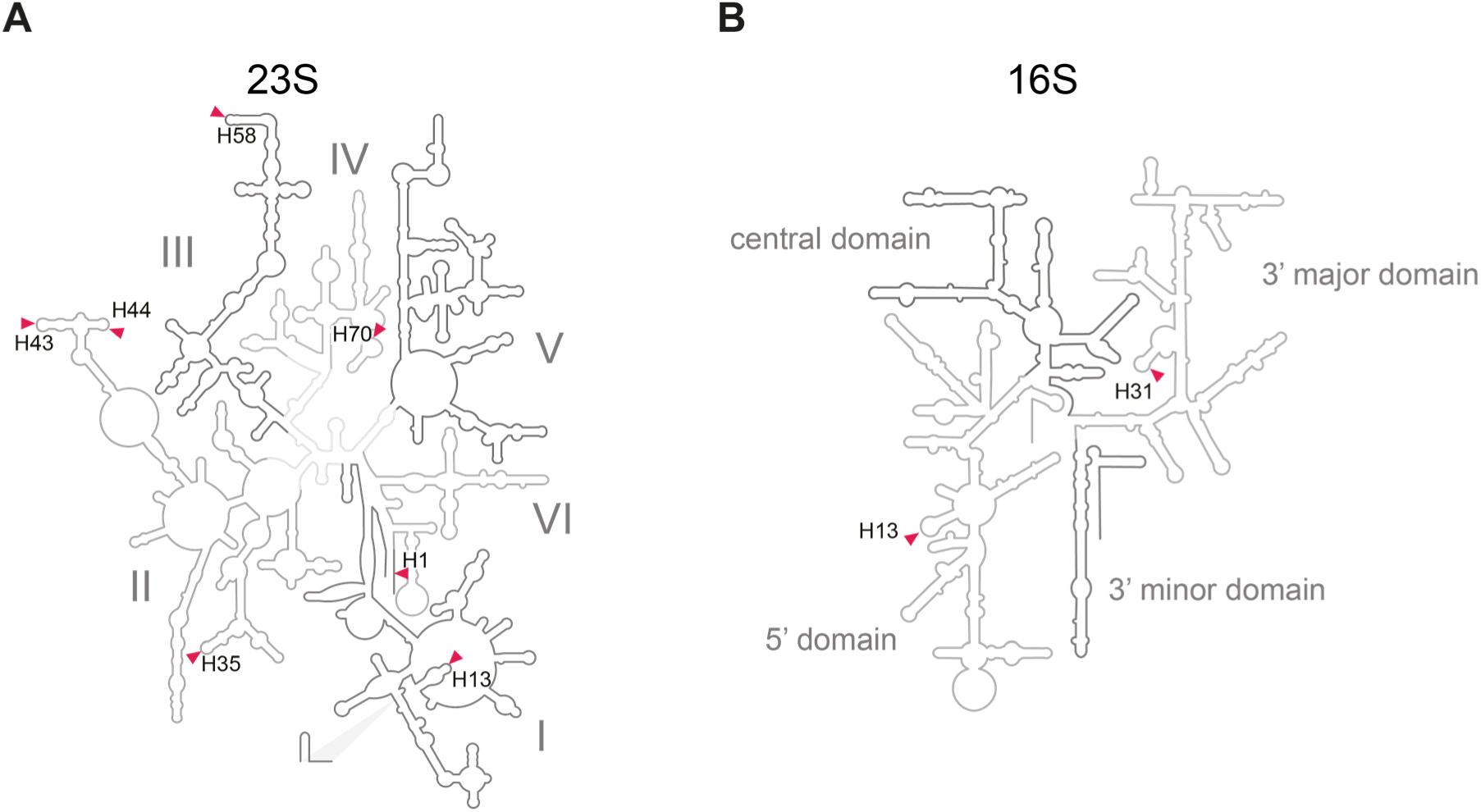
Mapping of Y-complex-dependent processing in the secondary structure of rRNA at the late exponential phase. Cleavages are marked with an arrow and the helix number for **A)** 23S and **B)** 16S rRNA.

**Supplementary Figure 6.**
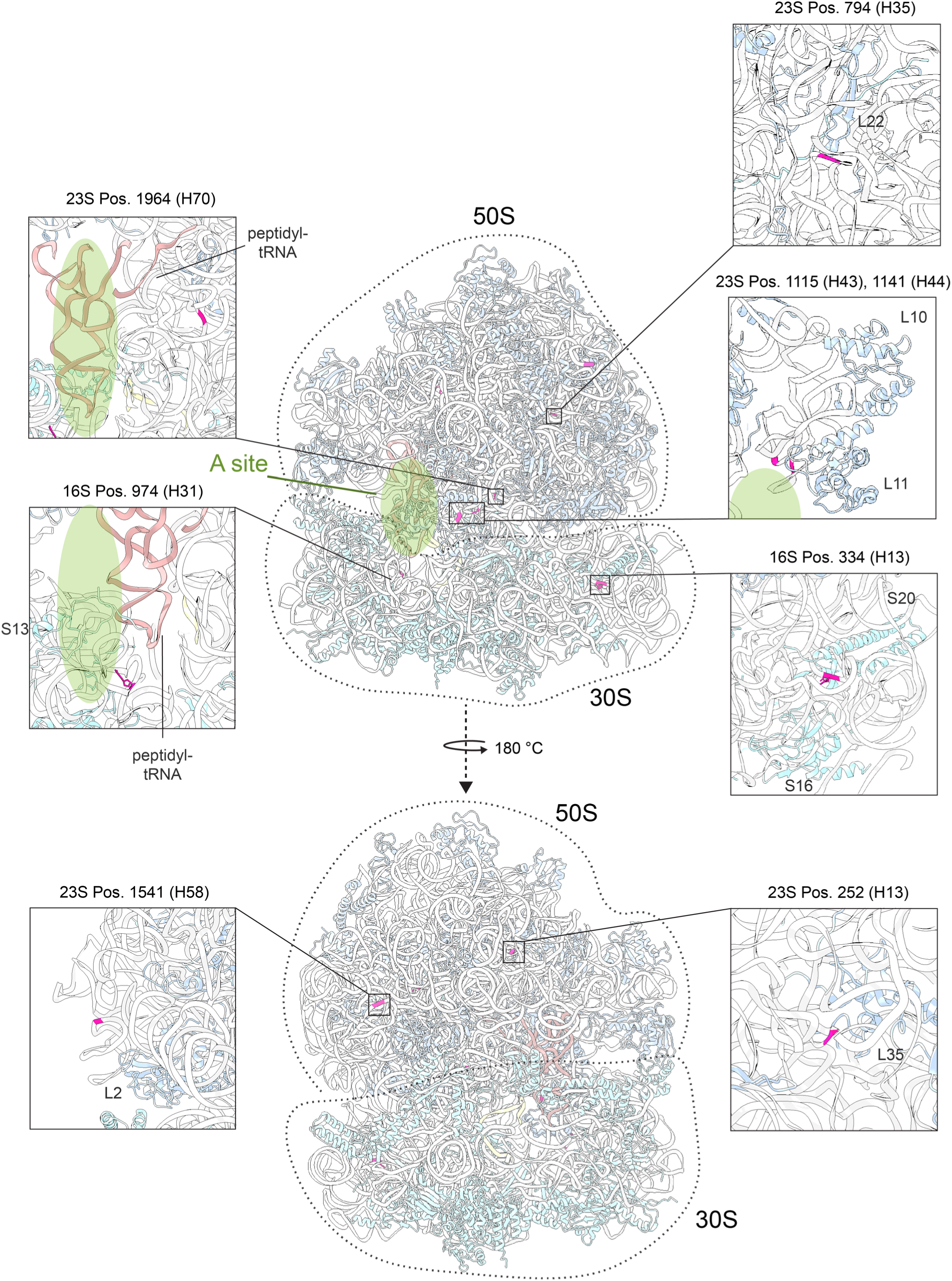
Mapping of Y-complex dependent processing in the ribosome 3D structure. The RNA end detected by the ISCP method is highlighted in magenta. The ribosome structure was retrieved from PDB (ID: 3J9W)^85^. The A site is shown as a green ellipse.

**Supplementary Figure 7.**
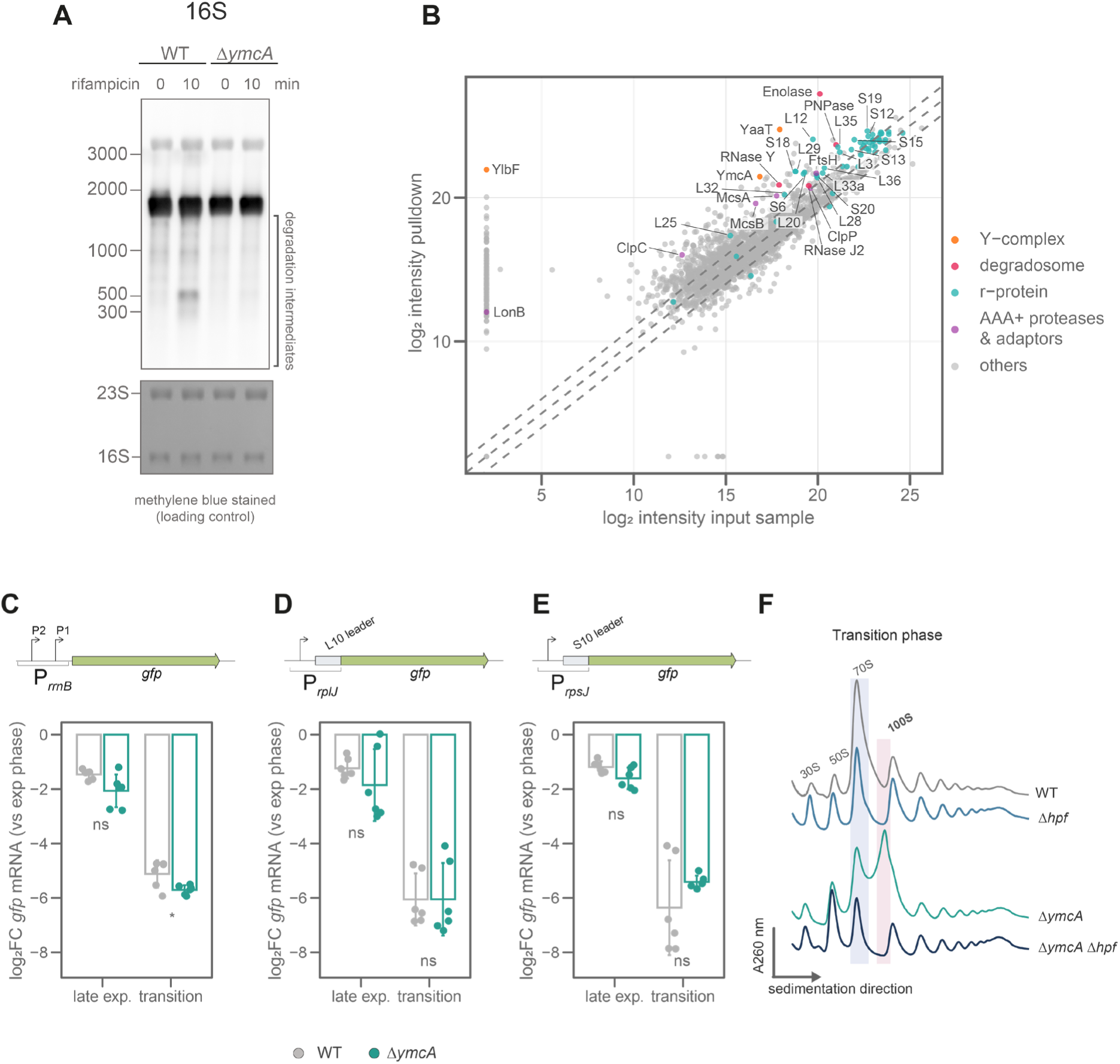
Y-complex-dependent initiates degradation of rRNA but does not affect transcriptional control of stringent response. **A)** Northern blot against 16S of WT and Δ*ymcA* cells at the late exponential phase, before and after treatment with 100 µg/ml rifampicin to inhibit transcription. The methylene blue-stained membrane is shown as a loading control. This image is representative of three biological replicates. **B)** Pulldown of YaaT-FLAG at the late exponential phase. The protein intensities of the input sample and the elution fraction were measured by mass spectrometry. The dotted lines indicate a fold change of less than 2. A log_2_ intensity value of two was inputed when a protein was missing in one of the samples. Activity of the promoter controlling **C)** *rrnB* (rRNA) transcription, **D)** *rplJ*, and **E)** *rpsJ* (r-proteins) genes at late exponential and transition phase compared to the exponential phase in WT and Δ*ymcA*. The promoters were cloned controlling *gfp* transcription, which was measured using RT-qPCR. The *pcp* transcript was used as a reference. Individual values and the average ± standard deviation of three biological replicates and two technical replicates are shown. The statistical significance was tested using student’s *t*-test. ns: *p-*value > 0.05, *: *p-*value ≤ 0.05. **F)** Ribosome sedimentation profile in 10-50% sucrose gradients of WT, Δ*hpf,* Δ*ymcA,* and Δ*ymcA* Δ*hpf* in the transition phase. The 70S and 100S ribosomes are highlighted with a blue or red box, respectively. The data is representative of three biological replicates.

**Supplementary Figure 8.**
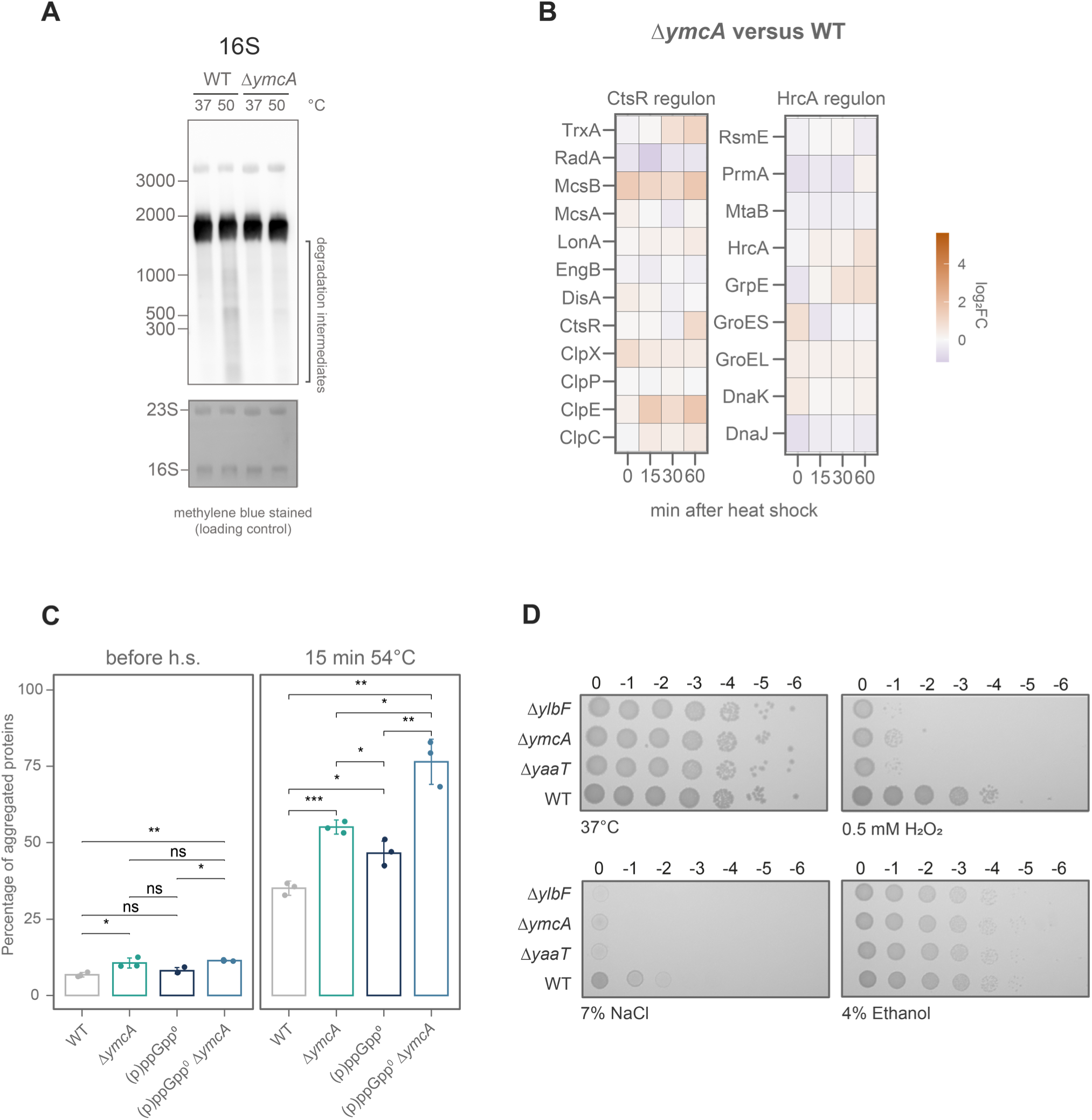
Control of ribosome levels and protein homeostasis in *ymcA* is important for responding against proteotoxic stresses. **A)** Northern blot against 16S of WT and Δ*ymcA* cells at exponential phase before and after heat shock. The methylene blue-stained membrane is shown as a loading control. This image is representative of three biological replicates. **B)** Induction of CtsR and HrcA regulons after heat shock (50°C) Δ*ymcA* compared to the WT. Proteins were measured by mass spectrometry. The color bar shows the log_2_ fold change compared to the WT control. **C)** Percentage of aggregated proteins before and after a 15 min heat shock at 54°C in WT and Δ*ymcA*, (p)ppGpp^0^ and (p)ppGpp^0^ Δ*ymcA*. Individual values and the average ± standard deviation of three biological replicates are shown. The statistical difference was tested using student’s *t*-test. ns: *p-*value > 0.05, *: *p-*value ≤ 0.05, **: *p-*value ≤ 0.01, ***: *p-*value ≤ 0.001. **D)** Spot test of deletion mutants of Y-complex members in the displayed stresses. The picture is representative of three biological replicates.

## References

1. Hecker, M., Schumann, W. & Völker, U. Heat-shock and general stress response in *Bacillus subtilis*. Mol. Microbiol. 19, 417–428 (1996).

2. Mogk, A. et al. The GroE chaperonin machine is a major modulator of the CIRCE heat shock regulon of *Bacillus subtilis*. EMBO J. 16, 4579–4590 (1997).

3. Krüger, E. & Hecker, M. The first gene of the *Bacillus subtilis clpC* operon, *ctsR*, encodes a negative regulator of its own operon and other Class III heat shock genes. J. Bacteriol. 180, 6681–6688 (1998).

4. Elsholz, A. K. W., Michalik, S., Zühlke, D., Hecker, M. & Gerth, U. CtsR, the Gram-positive master regulator of protein quality control, feels the heat. EMBO J. 29, 3621–3629 (2010).

5. Krüger, E., Witt, E., Ohlmeier, S., Hanschke, R. & Hecker, M. The Clp proteases of *Bacillus subtilis* are directly involved in degradation of misfolded proteins. J. Bacteriol. 182, 3259–3265 (2000).

6. Alver, R. et al. The control of protein arginine phosphorylation facilitates proteostasis by an AAA+ chaperone protease system. 2022.09.15.508104 Preprint at 10.1101/2022.09.15.508104 (2023).

7. Schäfer, H. et al. The alarmones (p)ppGpp are part of the heat shock response of *Bacillus subtilis*. PLOS Genet. 16, e1008275 (2020).

8. Clay, K. J., Yang, Y., Clark, C. & Petrascheck, M. Proteostasis is differentially modulated by inhibition of translation initiation or elongation. eLife 12, e76465 (2023).

9. Cherkasov, V. et al. Coordination of translational control and protein homeostasis during severe heat stress. Curr. Biol. 23, 2452–2462 (2013).

10. Schramm, F. D., Schroeder, K. & Jonas, K. Protein aggregation in bacteria. FEMS Microbiol. Rev. 44, 54–72 (2020).

11. Schäfer, H. et al. Spx, the central regulator of the heat and oxidative stress response in *B. subtilis*, can repress transcription of translation-related genes. Mol. Microbiol. mmi.14171 (2018) doi:10.1111/mmi.14171.

12. Nakano, S., Erwin, K. N., Ralle, M. & Zuber, P. Redox-sensitive transcriptional control by a thiol/disulphide switch in the global regulator, Spx. Mol. Microbiol. 55, 498–510 (2005).

13. Runde, S. et al. The role of thiol oxidative stress response in heat-induced protein aggregate formation during thermotolerance in *Bacillus subtilis*. Mol. Microbiol. 91, 1036–1052 (2014).

14. Rochat, T. et al. Genome-wide identification of genes directly regulated by the pleiotropic transcription factor Spx in *Bacillus subtilis*. Nucleic Acids Res. 40, 9571–9583 (2012).

15. Diez, S., Ryu, J., Caban, K., Gonzalez, R. L. & Dworkin, J. The alarmones (p)ppGpp directly regulate translation initiation during entry into quiescence. Proc. Natl. Acad. Sci. 117, 15565–15572 (2020).

16. Corrigan, R. M., Bellows, L. E., Wood, A. & Gründling, A. ppGpp negatively impacts ribosome assembly affecting growth and antimicrobial tolerance in Gram-positive bacteria. Proc. Natl. Acad. Sci. 113, (2016).

17. Pausch, P. et al. Structural basis for (p)ppGpp-mediated inhibition of the GTPase RbgA. J. Biol. Chem. 293, 19699–19709 (2018).

18. Krásný, L., Tišerová, H., Jonák, J., Rejman, D. & Šanderová, H. The identity of the transcription +1 position is crucial for changes in gene expression in response to amino acid starvation in *Bacillus subtilis*. Mol. Microbiol. 69, 42–54 (2008).

19. Krásný, L. & Gourse, R. L. An alternative strategy for bacterial ribosome synthesis: *Bacillus subtilis* rRNA transcription regulation. EMBO J. 23, 4473–4483 (2004).

20. Koo, B.-M. et al. Construction and analysis of two genome-scale deletion libraries for *Bacillus subtilis*. Cell Syst. 4, 291–305.e7 (2017).

21. Msadek, T. et al. ClpP of *Bacillus subtilis* is required for competence development, motility, degradative enzyme synthesis, growth at high temperature and sporulation. Mol. Microbiol. 27, 899–914 (1998).

22. Hajdusits, B. et al. McsB forms a gated kinase chamber to mark aberrant bacterial proteins for degradation. eLife 10, e63505 (2021).

23. Matavacas, J., Hallgren, J. & von Wachenfeldt, C. *Bacillus subtilis* forms twisted cells with cell wall integrity defects upon removal of the molecular chaperones DnaK and trigger factor. Front. Microbiol. 13, (2023).

24. Schumann, W. Regulation of bacterial heat shock stimulons. Cell Stress Chaperones 21, 959–968 (2016).

25. DeLoughery, A., Dengler, V., Chai, Y. & Losick, R. Biofilm formation by *Bacillus subtilis* requires an endoribonuclease-containing multisubunit complex that controls mRNA levels for the matrix gene repressor SinR. Mol. Microbiol. 99, 425–437 (2016).

26. DeLoughery, A., Lalanne, J.-B., Losick, R. & Li, G.-W. Maturation of polycistronic mRNAs by the endoribonuclease RNase Y and its associated Y-complex in *Bacillus subtilis*. Proc. Natl. Acad. Sci. 115, E5585–E5594 (2018).

27. Broglia, L. et al. An RNA-seq based comparative approach reveals the transcriptome-wide interplay between 3′-to-5′ exoRNases and RNase Y. Nat. Commun. 11, 1587 (2020).

28. Taggart, J. C. et al. A high-resolution view of RNA endonuclease cleavage in *Bacillus subtilis*. Nucleic Acids Res. 53, gkaf030 (2025).

29. Khemici, V., Prados, J., Linder, P. & Redder, P. Decay-initiating endoribonucleolytic cleavage by RNase Y is kept under tight control via sequence preference and sub-cellular localisation. PLOS Genet. 11, e1005577 (2015).

30. Le Scornet, A. et al. Critical factors for precise and efficient RNA cleavage by RNase Y in *Staphylococcus aureus*. PLOS Genet. 20, e1011349 (2024).

31. Shahbabian, K., Jamalli, A., Zig, L. & Putzer, H. RNase Y, a novel endoribonuclease, initiates riboswitch turnover in *Bacillus subtilis*. EMBO J. 28, 3523–3533 (2009).

32. Chao, Y. et al. In vivo cleavage map illuminates the central role of RNase E in coding and non-coding RNA pathways. Mol. Cell 65, 39–51 (2017).

33. Kime, L., Jourdan, S. S., Stead, J. A., Hidalgo-Sastre, A. & McDowall, K. J. Rapid cleavage of RNA by RNase E in the absence of 5′ monophosphate stimulation. Mol. Microbiol. 76, 590–604 (2010).

34. Davis, J. H. & Williamson, J. R. Structure and dynamics of bacterial ribosome biogenesis. Philos. Trans. R. Soc. B Biol. Sci. 372, 20160181 (2017).

35. Dubnau, E., DeSantis, M. & Dubnau, D. Formation of a stable RNase Y-RicT (YaaT) complex requires RicA (YmcA) and RicF (YlbF). mBio e01269–23 (2023) doi:10.1128/mbio.01269-23.

36. Eymann, C., Homuth, G., Scharf, C. & Hecker, M. *Bacillus subtilis* functional genomics: global characterization of the stringent response by proteome and transcriptome analysis. J. Bacteriol. 184, 2500–2520 (2002).

37. Tagami, K. et al. Expression of a small (p)ppGpp synthetase, YwaC, in the (p)ppGpp^0^ mutant of *Bacillus subtilis* triggers YvyD-dependent dimerization of ribosome. MicrobiologyOpen 1, 115–134 (2012).

38. Beckert, B. et al. Structure of the *Bacillus subtilis* hibernating 100S ribosome reveals the basis for 70S dimerization. EMBO J. 36, 2061–2072 (2017).

39. Carabetta, V. J. et al. A complex of YlbF, YmcA and YaaT regulates sporulation, competence and biofilm formation by accelerating the phosphorylation of Spo0A. Mol. Microbiol. 88, 283–300 (2013).

40. Tortosa, P., Albano, M. & Dubnau, D. Characterization of *ylbF*, a new gene involved in competence development and sporulation in *Bacillus subtilis*. Mol. Microbiol. 35, 1110–1119 (2000).

41. Branda, S. S. et al. Genes involved in formation of structured multicellular communities by *Bacillus subtilis*. J. Bacteriol. 186, 3970–3979 (2004).

42. Segev, E., Smith, Y. & Ben-Yehuda, S. RNA dynamics in aging bacterial spores. Cell 148, 139–149 (2012).

43. Hinrichs, R., Pozhydaieva, N., Höfer, K. & Graumann, P. L. Y-complex proteins show RNA-dependent binding events at the cell membrane and distinct single-molecule dynamics. Cells 11, 933 (2022).

44. Adusei-Danso, F. et al. Structure-function studies of the *Bacillus subtilis* Ric proteins identify the Fe-S cluster-ligating residues and their roles in development and RNA processing. mBio 10, e01841–19, /mbio/10/5/mBio.01841-19.atom (2019).

45. Lehnik-Habrink, M., et al. The RNA degradosome in *Bacillus subtilis*: identification of CshA as the major RNA helicase in the multiprotein complex. Mol. Microbiol. 77, 958–971 (2010).

46. Commichau, F. M. et al. Novel activities of glycolytic enzymes in *Bacillus subtilis*. Mol. Cell. Proteomics 8, 1350–1360 (2009).

47. Durand, S. & Condon, C. RNases and helicases in Gram-positive bacteria. Microbiol. Spectr. 6, 10.1128/microbiolspec.rwr-0003–2017.

48. Dimitrova-Paternoga, L. et al. Structural basis of ribosomal 30S subunit degradation by RNase R. Nature 626, 1133–1140 (2024).

49. Boylan, S. A., Redfield, A. R., Brody, M. S. & Price, C. W. Stress-induced activation of the Sigma B transcription factor of *Bacillus subtilis*. J. Bacteriol. 175, 7931–7937 (1993).

50. Kirstein, J., Molière, N., Dougan, D. A. & Turgay, K. Adapting the machine: adaptor proteins for Hsp100/Clp and AAA+ proteases. Nat. Rev. Microbiol. 7, 589–599 (2009).

51. Lehnik-Habrink, M. et al. RNA processing in *Bacillus subtilis*: identification of targets of the essential RNase Y: Targets of the essential RNase Y in *Bacillus subtilis*. Mol. Microbiol. 81, 1459–1473 (2011).

52. Anderson, B. W., Fung, D. K. & Wang, J. D. Regulatory themes and variations by the stress-signaling nucleotide alarmones (p)ppGpp in bacteria. Annu. Rev. Genet. 55, 115–133 (2021).

53. Driller, K., Cornejo, F. A. & Turgay, K. (p)ppGpp – an important player during heat shock response. microLife 4, uqad017 (2023).

54. Zundel, M. A., Basturea, G. N. & Deutscher, M. P. Initiation of ribosome degradation during starvation in *Escherichia coli*. RNA 15, 977–983 (2009).

55. Sulthana, S., Basturea, G. N. & Deutscher, M. P. Elucidation of pathways of ribosomal RNA degradation: an essential role for RNase E. RNA 22, 1163–1171 (2016).

56. Lehnik-Habrink, M. et al. RNase Y in *Bacillus subtilis*: a natively disordered protein that is the functional equivalent of RNase E from *Escherichia coli*. J. Bacteriol. 193, 5431–5441 (2011).

57. Marcaida, M. J., DePristo, M. A., Chandran, V., Carpousis, A. J. & Luisi, B. F. The RNA degradosome: life in the fast lane of adaptive molecular evolution. Trends Biochem. Sci. 31, 359–365 (2006).

58. Piir, K., Paier, A., Liiv, A., Tenson, T. & Maiväli, Ü. Ribosome degradation in growing bacteria. EMBO Rep. 12, 458–462 (2011).

59. Kuroda, A. et al. Role of inorganic polyphosphate in promoting ribosomal protein degradation by the Lon protease in *E. coli*. Science 293, 705–708 (2001).

60. An, H. & Harper, J. W. Ribosome abundance control via the ubiquitin–proteasome system and autophagy. J. Mol. Biol. 432, 170–184 (2020).

61. Huang, Z. et al. RIOK3 mediates the degradation of 40S ribosomes. Mol. Cell 85, 802–814.e12 (2025).

62. Ford, P. W. et al. RNF10 and RIOK3 facilitate 40S ribosomal subunit degradation upon 60S biogenesis disruption or amino acid starvation. Cell Rep. 44, 115371 (2025).

63. Coria, A. R. et al. The integrated stress response regulates 18S nonfunctional rRNA decay in mammals. Mol. Cell 85, 787–801.e8 (2025).

64. Simpkins, S. W. et al. Using BEAN-counter to quantify genetic interactions from multiplexed barcode sequencing experiments. Nat. Protoc. 14, 415–440 (2019).

65. Benjamini, Y. & Hochberg, Y. Controlling the false discovery rate: a practical and powerful approach to multiple testing. J. R. Stat. Soc. Ser. B Methodol. 57, 289–300 (1995).

66. Lécrivain, A.-L. et al. In vivo 3′-to-5′ exoribonuclease targetomes of *Streptococcus pyogenes*. Proc. Natl. Acad. Sci. 115, 11814–11819 (2018).

67. Martin, M. Cutadapt removes adapter sequences from high-throughput sequencing reads. EMBnet.journal 17, 10 (2011).

68. Dobin, A. et al. STAR: ultrafast universal RNA-seq aligner. Bioinformatics 29, 15–21 (2013).

69. Li, H. et al. The Sequence Alignment/Map format and SAMtools. Bioinformatics 25, 2078–2079 (2009).

70. Smith, T., Heger, A. & Sudbery, I. UMI-tools: modeling sequencing errors in Unique Molecular Identifiers to improve quantification accuracy. Genome Res. 27, 491–499 (2017).

71. Liao, Y., Smyth, G. K. & Shi, W. featureCounts: an efficient general purpose program for assigning sequence reads to genomic features. Bioinformatics 30, 923–930 (2014).

72. Love, M. I., Huber, W. & Anders, S. Moderated estimation of fold change and dispersion for RNA-seq data with DESeq2. Genome Biol. 15, 550 (2014).

73. Putri, G. H., Anders, S., Pyl, P. T., Pimanda, J. E. & Zanini, F. Analysing high-throughput sequencing data in Python with HTSeq 2.0. Bioinformatics 38, 2943–2945 (2022).

74. Mölder, F., et al. Sustainable data analysis with Snakemake. Preprint at 10.12688/f1000research.29032.2 (2021).

75. McCarthy, D. J., Chen, Y. & Smyth, G. K. Differential expression analysis of multifactor RNA-Seq experiments with respect to biological variation. Nucleic Acids Res. 40, 4288–4297 (2012).

76. Geissler, A. S. et al. BSGatlas: a unified *Bacillus subtilis* genome and transcriptome annotation atlas with enhanced information access. Microb. Genomics 7, 000524 (2021).

77. Tareen, A. & Kinney, J. B. Logomaker: beautiful sequence logos in Python. Bioinformatics 36, 2272–2274 (2020).

78. Lorenz, R. et al. ViennaRNA Package 2.0. Algorithms Mol. Biol. 6, 26 (2011).

79. Elfmann, C., Dumann, V., van den Berg, T. & Stülke, J. A new framework for SubtiWiki, the database for the model organism *Bacillus subtilis*. Nucleic Acids Res. 53, D864–D870 (2025).

80. Pfaffl, M. W. A new mathematical model for relative quantification in real-time RT-PCR. Nucleic Acids Res. 29, 45e–445 (2001).

81. Hughes, C. S. et al. Single-pot, solid-phase-enhanced sample preparation for proteomics experiments. Nat. Protoc. 14, 68–85 (2019).

82. Kohler, D. et al. MSstats version 4.0: statistical analyses of quantitative mass spectrometry-based proteomic experiments with chromatography-based quantification at scale. J. Proteome Res. 22, 1466–1482 (2023).

83. Jahn, M. et al. Growth of cyanobacteria is constrained by the abundance of light and carbon assimilation proteins. Cell Rep. 25, 478–486.e8 (2018).

84. Jahn, M. et al. Protein allocation and utilization in the versatile chemolithoautotroph *Cupriavidus necator*. eLife 10, e69019 (2021).

85. Sohmen, D. et al. Structure of the *Bacillus subtilis* 70S ribosome reveals the basis for species-specific stalling. Nat. Commun. 6, 6941 (2015).

